# CLAVATA signalling shapes barley inflorescence architecture by controlling activity and determinacy of shoot apical and rachilla meristems

**DOI:** 10.1101/2024.05.28.595952

**Authors:** Isaia Vardanega, Jan Eric Maika, Edgar Demesa-Arevalo, Tianyu Lan, Gwendolyn K. Kirschner, Jafargholi Imani, Ivan F. Acosta, Katarzyna Makowska, Götz Hensel, Thilanka Ranaweera, Shin-Han Shiu, Thorsten Schnurbusch, Maria von Korff Schmising, Rüdiger Simon

## Abstract

Grasses exhibit a large variety of diverse inflorescence architectures, from complex branched inflorescences in *Oryzeae* (rice) to simple spike-type inflorescences in *Triticeae* (e.g. barley, wheat). Inflorescence architecture depends on shape, longevity and determinacy of meristems that direct growth of the main rachis and lateral branches, but how individual meristem activities are determined and integrated within complex inflorescences is not yet understood. We found that activity of distinct meristems in the barley inflorescence is coordinated by a signalling pathway comprising the receptor like kinase *Hordeum vulgare* CLAVATA1 (HvCLV1) and the secreted CLAVATA3/ENDOSPERM SURROUNDING REGION (CLE)-family peptide FON2- LIKE CLE PROTEIN1 (HvFCP1). HvFCP1 interacts with HvCLV1 to promote spikelet formation but restricts inflorescence meristem and rachilla meristem proliferation. *Hvfcp1* or *Hvclv1* mutants generate branched inflorescences with additional rows of spikelets and supernumerary florets. Transcriptome analysis reveals that *HvFCP1/HvCLV1* signalling controls inflorescence branching through the regulation of trehalose-6-phosphate synthesis and sugar transport. Our discoveries reveal the potential to engineer barley inflorescence architecture by manipulating regulation of distinct meristem activities.

## Main

The *Poaceae* family (*Gramineae* or grasses) displays a large variety of inflorescence architectures that evolved from an ancestral compound spike with panicle-like branches on a main inflorescence axis (rachis)^1^. The multiple branching orders characteristic of panicles gradually simplified during evolution and domestication, generating the variety of current grass inflorescences^2^. The morphology of distinct grass inflorescence architectures depends on the initiation and placement of new meristems on the flanks of the shoot apical meristem (SAM), and on meristem size, longevity, identity and determinacy^3^. During the vegetative phase, the cereal SAM generates only leaf primordia in a distichous pattern. In response to both external and internal signals, the SAM converts into an inflorescence meristem (IM), which can initiate spikelets directly on the rachis in *Triticeae*^2^, including barley (*Hordeum vulgare* L.) and wheat (*Triticum* ssp.), or form primary and secondary branches^4^ in *Oryzeae* (rice) and *Andropogoneae* (maize and sorghum).

Spikelet meristems (SM) first give rise to two small modified bracts (the glumes), and later to a variable number of floret(s) that develop from floret meristems (FMs) on a short axis called rachilla, which is sustained by a rachilla meristem (RM). Florets carry the leaf-like lemma and palea, which enclose modified petals called lodicules, and the sex organs, the stamen and carpel^5^.

In barley, the IM remains indeterminate and generates triple spikelet meristems (TSM) on its flanks until developmentally programmed pre-anthesis tip degeneration causes IM senescence and death^5,6^. TSM splits into a central spikelet meristem (CSM), flanked by two lateral spikelet meristems (LSM). In two-rowed cultivars, lateral spikelets remain sterile and arrest before floret organs are fully developed^7^. Each SM further divides into the vestigial rachilla meristem (RM), an abaxial floret meristem (FM), and a subtending lemma primordium (LEP).

Inflorescence branching in barley is suppressed by the transcription factors COMPOSITUM1 (COM1), COM2^8,9^ and HvMADS1, while INTERMEDIUM-m restricts floret number per spikelet and maintains indeterminacy of the IM^10,11^. In maize, inflorescence branching is controlled by the RAMOSA pathway, comprising RAMOSA3 (RA3), a Trehalose-6-Phosphate Phosphatase that putatively regulates branching by a sugar signal that moves into axillary meristems^12^. Importantly, even closely related grasses, such as the temperate cereals wheat and barley, differ in their inflorescence architecture because of differences in meristem behaviour. In wheat (*Triticum aestivum* L.), inflorescence growth arrests with differentiation of the IM into an SM, but the RM remains indeterminate, enabling the formation of up to 12 florets^2,5^. These examples illustrate how differential regulation of meristem activities finally impact inflorescence architecture. However, the regulatory networks, feedback regulations and external inputs that coordinate the size, longevity and determinacy of different meristem types in the inflorescence are still unknown.

In Arabidopsis, activity of shoot and floral meristems depends on the WUSCHEL transcription factor, which moves from a deeper meristem region to the stem cell zone to promote stem cell maintenance by suppressing auxin response factors. WUSCHEL promotes expression of the CLAVATA3/ENDOSPERM SURROUNDING REGION (CLE) family peptide CLV3, which is secreted from stem cells and interacts with leucine-rich-repeat receptor kinases (LRR-RLKs) of the CLAVATA1 (CLV1) family to repress WUSCHEL, thereby providing a negative feedback signal. CLE40 acts from the meristem periphery through the CLV1-related RLK BARELY ANY MERISTEM 1 (BAM1) to impact meristem shape^13^. CLV-related signalling pathways regulate diverse meristem activities, including root and cambial meristems^14^, and CLE / CLV1-family signalling has also been described for rice and maize. In rice, the LRR-RLK FLORAL ORGAN NUMBER1 (FON1) and the CLE peptide FLORAL ORGAN NUMBER2 (FON2) restrict the sizes of FMs, IM and the number of primary branches^15,16^, while FON2-LIKE CLE PROTEIN1 (FCP1), which is highly conserved between cereal grasses, likely plays an antagonistic role and promotes maintenance of the vegetative SAM and root apical meristem^17,18^. The maize LRR-RLK THICK TASSEL DWARF1 (TD1) and CLE7 confine the diameter of both tassel and ear meristems. In *td1* mutants, an enlarged IM initiates disorganized supernumerary rows of spikelet pair meristems, which sometimes develop additional SMs^19,20^. In parallel, ZmFCP1 signalling suppresses stem cell proliferation in the ear meristem, which is enlarged in *Zmfcp1* mutants^18,20^.

The development of cereal inflorescence architectures requires a close coordination of meristem ontogenies. Here, we investigated how CLV-related signalling pathways may contribute to this process in barley. We identify the barley HvCLV1 and HvFCP1 and show that they act in joint, but also separate, signalling pathways to impact multiple aspects of meristem development. Our findings extend the roles of CLV signalling, from feedback signalling in stem cell homeostasis, to coordination of meristem shape, organ formation and determinacy in cereal inflorescences.

## Results

To identify CLV1-related RLKs from barley, we analysed the phylogeny for all protein kinase sequences from two dicotyledons (*Arabidopsis thaliana*, *Solanum lycopersicum*) and four gramineous species belonging to the *Poaceae* family (*Zea mays*, *Oryza sativa japonica*, *Triticum turgidum* and *Hordeum vulgare*) (Suppl.Data1). Within the clade comprising AtCLV1, we identified six closely related genes from barley. HORVU.MOREX.r3.7HG0747230 represented the closest ortholog of AtCLV1 in *Hordeum vulgare* and was named *HvCLV1*. *HvCLV1* grouped with the maize and rice orthologs *ZmTD1* and *OsFON1*, the other five genes in the clade were more closely related to *AtBAM1 to 3* and we designated them *HvBAM1* to *5* (ExtDataFig.1A). *HvCLV1* encodes an LRR-RLK protein of 1,015 amino acids, comprising an intracellular kinase domain and 20 extracellular Leucin Rich Repeats (LRRs), similar to the closely related ZmTD1 and OsFON1, and AtCLV1 with 21 LRRs (Fig.1A). *HvCLV1* expression was analysed using single-molecule RNA fluorescent *in situ* hybridisation (smRNA-FISH, Molecular Cartography^TM^, Resolve Biosciences) on sectioned developing barley apices during vegetative (Fig.1B-D) and reproductive stages (Fig.1E-M, ExtDataFig.1B), and HvCLV1 protein localization was analysed using the translational reporter line *pHvCLV1:HvCLV1- mVenus*, which expresses the HvCLV1 protein with the fluorophore mVenus fused C-terminally to the cytoplasmic kinase domain and functionally complements a *Hvclv1* mutant (see below, ExtDataFig.2N). We used the Waddington scale (Waddington stage, W) to define stages of barley development^21^. During vegetative (W1) and reproductive development (W3.5) (Fig.1B,E), *HvCLV1* is expressed mostly in the three outer cell layers in the apical meristem (Fig.1C,F), and throughout spikelet development in RM, FM, lemma primordium and flower organs (Fig.1H,L). *HvCLV1* mRNA and HvCLV1 protein were detected in similar patterns (Fig.1D,G,I,M). Longitudinal sectioning of the developing spikelet showed HvCLV1 internalization in the RM (Fig.1J,K, ExtDataFig.1D). HvCLV1 localised to the plasma membrane and to cytoplasmic structures, which could reflect either *de novo* synthesis and intracellular trafficking, or turnover after signalling^22^.

**Fig. 1:**
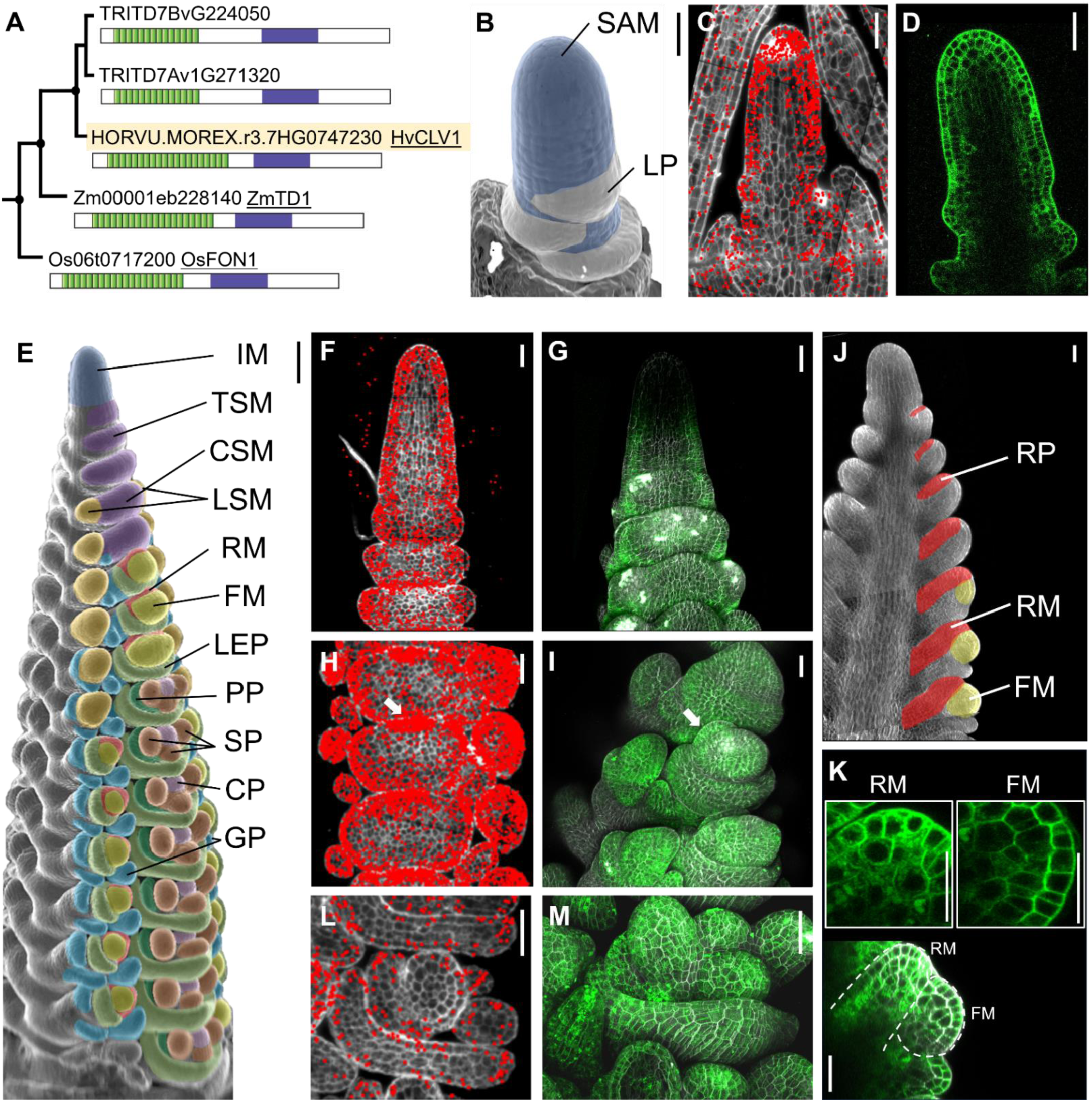
Identification and expression pattern of HvCLV1 in meristems composing the barley inflorescence. **(A)** Maximum likelihood tree of HvCLV1 subclade. Dots indicate nodes with bootstrap value higher than 80. Gene identifiers are shown next to a schematic representation of protein structures. Kinase domain in purple and LRRs in green. HvCLV1 highlighted in light orange. **(B)** SEM picture of barley vegetative meristem. Color code: shoot apical meristem meristem (SAM) in blue, leaf primordia (LP) in white. **(C)** smRNAfish detection of *HvCLV1* transcripts (red dots) at the vegetative stage, calcofluor stained cell wall in grey. **(D)** HvCLV1 protein localization in a central longitudinal section of the SAM at vegetative stage, HvCLV1 proteins tagged with mVenus in green**. (E)** *Hordeum vulgare* inflorescence cv. Golden Promise Fast, W3.5. Color code: inflorescence meristem (IM) in blue, triple spikelet meristem (TSM) and central spikelet meristem (CSM) in purple, lateral spikelet meristem (LSM) in orange, rachilla meristem (RM) in red, flower meristem (FM) in yellow, lemma primordia (LEP) in light green, palea primordia (PP) in dark green, stamen primordia (SP) in brown, carpel primordium (CP) in pink and glumes primordia (GP) in cyan. **(F,G)** Transcripts and proteins localization of HvCLV1 in the IM and TSMs, **(H,I)** in spikelet primordia at the FM initiation stage. The white arrows indicate the RM **(J)** Schematic representation of rachilla development. The rachilla primordium (RP) become RM after formation of the FM. **(K)** central longitudinal section of SM. Segmented lines indicate RM and FM. The close-up pictures show HvCLV1 proteins internalized in the vacuole in the RM (top left) and HvCLV1 proteins localized on the plasma membrane in the FM (top right). **(L,M)** HvCLV1 transcripts and proteins localization in stamens and carpel primordia. *HvCLV1* transcripts in red and HvCLV1 proteins in green. Scale bar = 50 µm, in (E) = 100 µm.

**Fig. 2:**
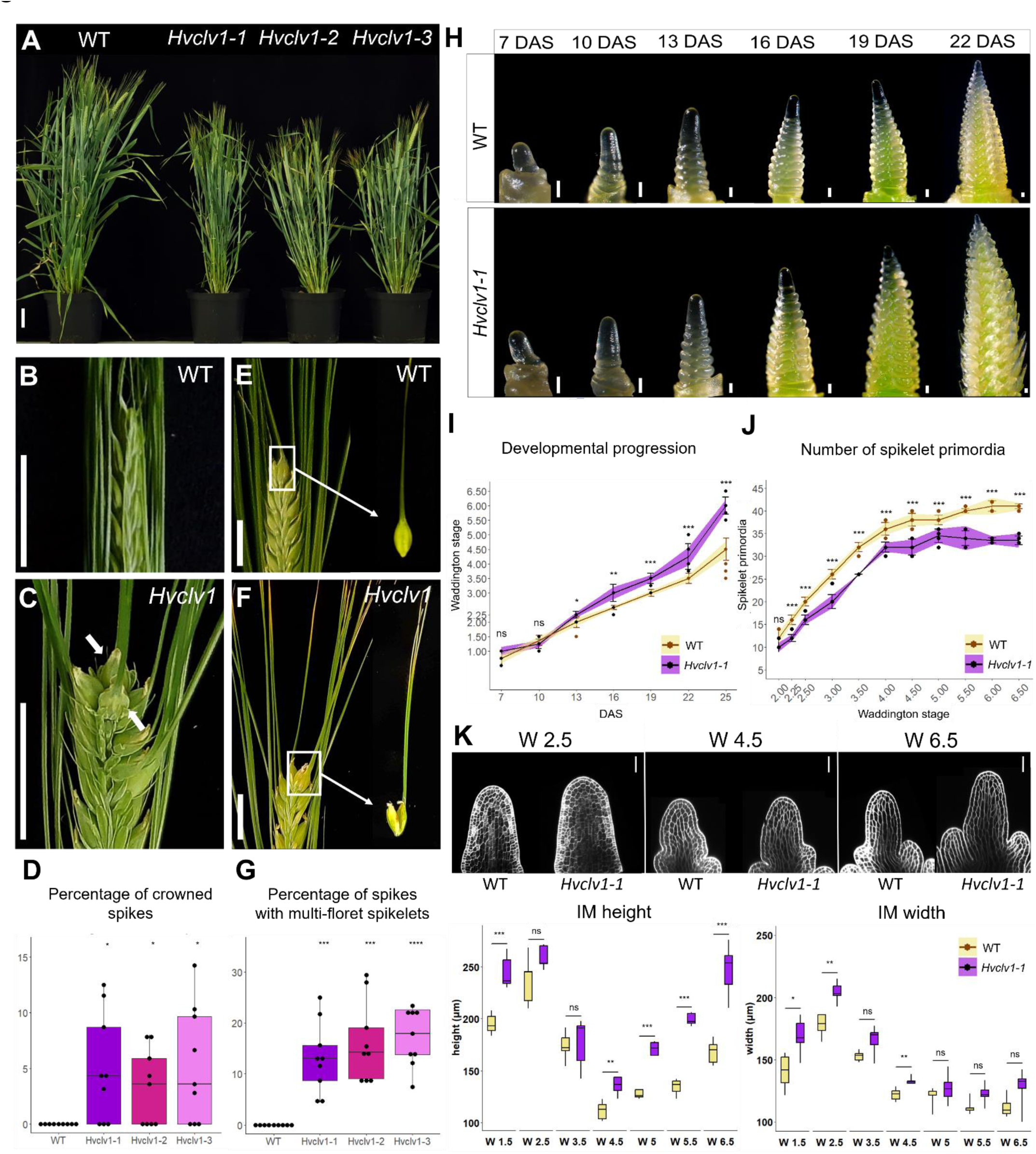
HvCLV1 plays a role in plant and spike architecture, delays inflorescence development and promotes spikelet formation. **(A)** Mature plants of Hordeum vulgare cv. Golden Promise Fast (WT) versus three selected *Hvclv1* mutant alleles (*Hvclv1-1*, *Hvclv1-2*, *Hvclv1-3*) **(B, C)** WT inflorescence and *Hvclv1-1* crowned spike phenotype respectively, ectopic grains are indicated by white arrows. **(D)** Percentage of crowned spikes in WT and *Hvclv1* mutant alleles. Dots represents the percentage per plant and asterixis indicate the significant difference to WT. n=9 **(E,F)** Close up respectively on WT single grain and *Hvclv1-1* multi-grain developed from multi-floret spikelets. **(G)** Percentage of spikes with multi-grain in WT plants and *Hvclv1* mutant alleles. Dots represents the percentage per plant and asterixis indicate the significant difference to WT. n=9 **(H)** Stereo microscope pictures of WT and *Hvclv1-1* SAM development from 7 to 22 days after sowing (DAS). **(I,J)** Developmental progression and number of spikelet primordia. WT in yellow, *Hvclv1-1* in purple. Dots represents single measurements; error bars represent standard deviation and the coloured ribbon the interval of confidence. n=10 (I) n =7 (J). **(K)** On the top, examples of WT and *Hvclv1-1* IM at W2.5, 4.5 and 6.5. The pictures were taken by confocal microscope, in white the cell wall stained with renaissance blue. The two plots on the bottom were generated by measuring meristem tip height and width at different W. Five samples per genotype were measured for every W. **(I)** Scale bar = 5 cm in (A), 1.5 cm (B, C, E, F), 100 μm (H) and 50 μm (K).

For a better understanding of the function of barley HvCLV1, we generated *Hvclv1* mutants by CRISPR-Cas9. Three independent alleles, *Hvclv1-1* to *-3*, which likely represent loss-of-function mutants, showed closely related phenotypes (ExtDataFig.2A, Suppl.info1). All *Hvclv1* mutants were semi-dwarfs (Fig.2A), with shorter stems, spikes and fewer internodes and tillers than WT (ExtDataFig.2B-E). *Hvclv1* mutants also developed fewer and smaller grains than WT (ExtDataFig.2F-K). Furthermore, a variable proportion of the *Hvclv1-1* spikes formed additional, ectopic rows of spikelets in a non-distichous phyllotaxis (crowned spikes) (Fig.2B- D), or carried multi-floret spikelets with two or three florets, separate embryos, and endosperms enclosed by partially fused lemmas (Fig.2E-G). These phenotypes were also observed in *Hvclv1* mutants grown in semi-field-like conditions in Germany between March to the end of July 2023, but not in WT (Suppl.info2). Detailed microscopic analysis of early development showed that *Hvclv1-1* and WT meristems developed similarly along vegetative stages (Fig.2H,I), although *Hvclv1* SAMs were slightly enlarged (see below). At W1.5 (between 10 and 13 DAS), meristems started to produce spikelet primordia while cells continued to accumulate in the IM until W2.5. IM size was then reduced during rapid spikelet initiation (W2.5 to W4.5). After termination of spikelet formation at W5 to W5.5, the width of the IM remained unaltered, and cells started to accumulate along the vertical axis (Fig.2I,J,K). *Hvclv1* mutant IMs were larger than WT IMs at W1.5, and developed faster than WT, reaching each stage earlier (Fig.2H-K). IMs of *Hvclv1* always appeared more elongated than WT IMs (Fig.2K), but also arrested spikelet initiation earlier (Fig.2J). The length of the entire inflorescences was briefly reduced at early stages when *Hvclv1* mutants progressed rapidly through developmental stages, but did not differ significantly from W4 onwards (ExtDataFig.2M). We conclude that *HvCLV1* first acts to restrict meristem growth and developmental progression of spikelet primordia, and promotes spikelet initiation at later stages.

To track the origin of the multi-floret spikelets and ectopic spikelet rows in *Hvclv1* plants, we imaged developing inflorescences by scanning electron microscopy (SEM). Bases of *Hvclv1* IMs were enlarged at the initiation of spikelet formation (W1.5) (Fig.2K), which correlated with the formation of an additional row of SMs in later stages. IMs then shifted from a distichous to a spiral phyllotaxis (Fig.3A,B). In WT, the barley SM gives rise to the rachilla meristem (RM), which arrests development after initiation of a single floret (Fig.3C-E). SEM analysis showed that the RM continues to grow wider and for an extended time in *Hvclv1* (Fig.3F), forming either larger or additional florets, or even secondary RMs (Fig.3G-J). We conclude that *HvCLV1* is required to restrict RM activities to the formation of a single floret.

**Fig. 3:**
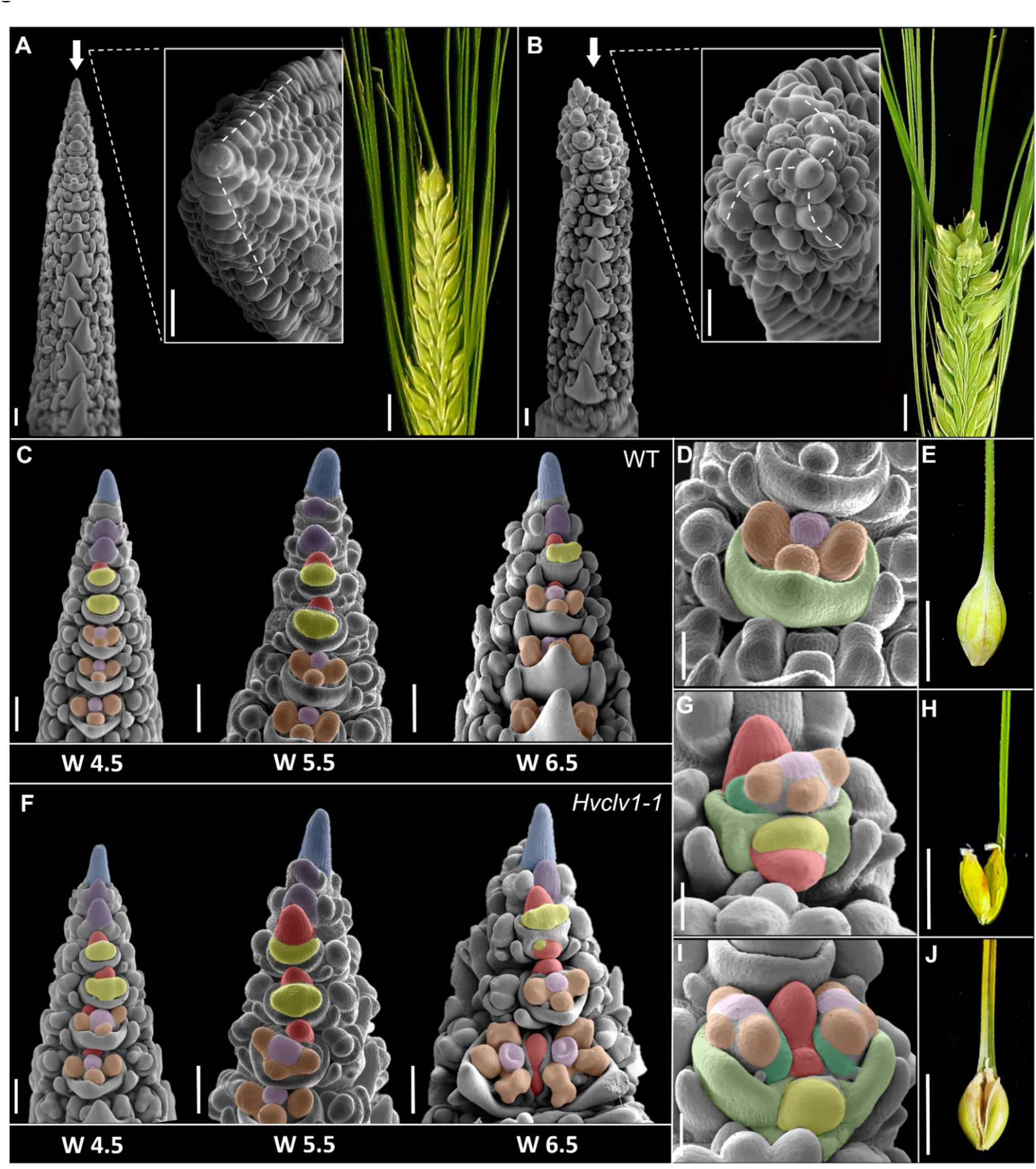
Origins of crowned spikes and multi-floret spikelets. **(A,B)** SEM pictures of WT and *Hvclv1-1* inflorescence meristems at W4.5 respectively. Frontal view on the left and close up of the top view in the center. Segmented lines indicate spikelet primordia phyllotaxis. On the right pictures of representative spike phenotypes. Crowned spikes were found in 6/38 *Hvclv1-1* inflorescences from W5.5 to W6.5 **(C)** Developmental progression of WT inflorescences at W4.5, 5.5 and 6.5. Color code as described in Fig.1E **(D,E)** Close up on WT floret and WT single grain respectively. **(F)** Developmental progression of *Hvclv1-1* inflorescences at W4.5, 5.5 and 6.5. Color code as described in Fig.1E. Multi-floret spikelets were found in 27/38 *Hvclv1-1* inflorescences from W5.5 to W6.5. **(G-J)** Close up on *Hvclv1-1* multi-floret spikelet and the resulting multi-grain disposed vertically (G,H) and horizontally (I,J). Color code (G-J): RM and secondary rachilla meristem (SRM) in red, FM in yellow, LP in light green, PP in dark green, SP in brown and CP in pink. Scale bars: A, B, C, F = 200 µm; D, G, I = 100 µm; E, H, J = 1 cm.

### The CLE peptide HvFCP1 acts with HvCLV1 to limit meristem activities

In many plant species, CLV1-family receptors were found to interact with CLE peptides closely related to Arabidopsis CLV3 and CLE40, such as OsFCP1 from rice. We found that the barley gene HORVU.MOREX.r3.2HG0174890 encodes an evolutionarily conserved FCP1-like peptide, which we named HvFCP1 (Suppl.info3). Incubation of WT barley seedlings with growth medium containing 30 µM of synthetic HvFCP1 peptide caused a reduction in meristem height of the WT SAM, while SAM width was not affected. *Hvclv1-1* mutant seedlings were insensitive to HvFCP1 treatment (Fig.4A), indicating that HvFCP1 requires HvCLV1 to limit SAM height.

**Fig. 4:**
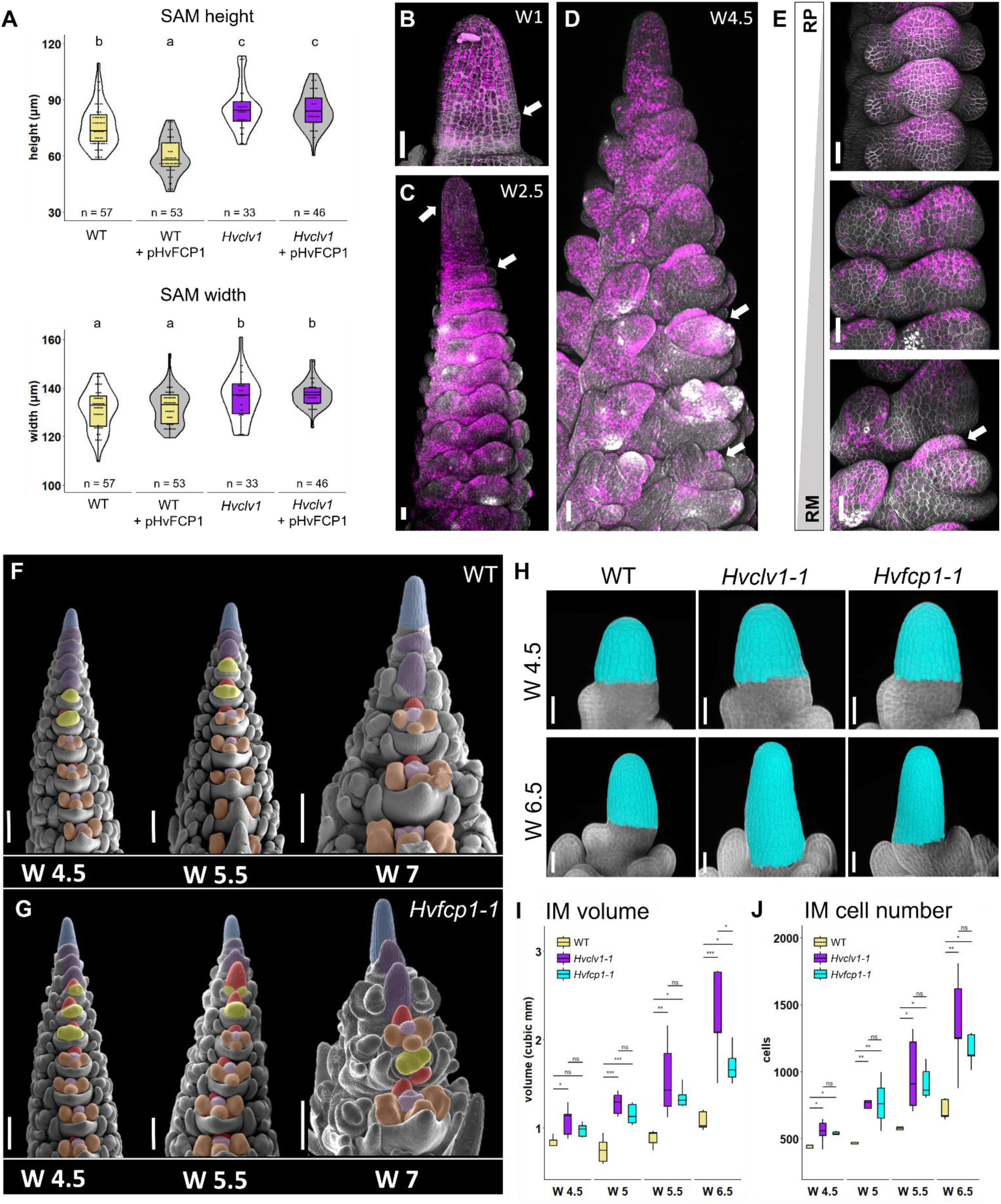
HvFCP1 interact with HvCLV1 to regulate IM proliferation and RM determinacy. **(A)** Vegetative SAM height and width in control samples (white) and samples treated with HvFCP1 synthetic peptide (pHvFCP1) (grey). WT SAMs in yellow and *Hvclv1-1* in purple. Dots represent single measurements, n = number of samples. Letters on top of each boxplot represent the results of the ANOVA test. **(B-D)** Confocal images of barley SAM and inflorescence expressing *HvFCP1* transcriptional reporter line (pHvFCP1:mVenus-H2B) at different developmental stages and organ primordia. SAM at vegetative stage (B), at W2.5 (C) and at W3.5 (D). **(E)** *HvFCP1* transcriptional reporter line along rachilla development. From rachilla primordium (RP) to rachilla meristem (RM). **(F,G)** Inflorescence phenotype in late stages of development (W4.5, W5.5, W7) in WT (F) and *Hvfcp1-1* (G). Multi-floret spikelets were found in 12/35 *Hvfcp1-1* inflorescences from W5.5 to W6.5. Color code as described in Fig.1E **(H-J)** 3D reconstruction of WT, *Hvclv1-1* and *Hvfcp1-1* IMs at W4.5 and W6.5 (H). Cells in cyan were selected for the IM measurements in (I) and (J), were boxplots displays IM volume and cell number respectively. WT (yellow), *Hvclv1-1* (purple), *Hvfcp1-1* (cyan). Scale bars: 50 µm; F and G = 200 µm.

We then analysed expression of a transcriptional reporter line, *pHvFCP1:mVenus-H2B*, in transgenic barley (Fig.4B-E). During the vegetative phase, *HvFCP1* promoter was active in the SAM, but downregulated in leaf initiation sites (Fig.4B). Later on, activity was found in the IM and from the triple spikelet meristem stage onwards (Fig.4C). Moreover, the activity was polarized at the adaxial side of the developing central spikelet, at the rachilla primordium (RP) and later in the fully formed RM (Fig.4E). In FMs, *HvFCP1* was mainly expressed on the central domain and later in carpel primordia (Fig.4D). Importantly, the *HvFCP1* reporter was more prominently expressed in the RM compared to *HvCLV1* (ExtDataFig.3A-J).

We generated two independent knock-out mutant alleles by CRISPR-Cas9 (*Hvfcp1-1* and *-2)* to study HvFCP1 function. Both *Hvfcp1* alleles are phenotypically indistinguishable, and molecular analysis identifies them as loss-of-function mutants (ExtDataFig.4A). *Hvfcp1* plants, similar to *Hvclv1*, remained shorter with shorter inflorescences and formed fewer viable grains, while tiller or internode number was not affected (ExtDataFig.4B-E). *Hvfcp1* mutant IM height and width increased similar to those of *Hvclv1* mutants, and developed faster than WT (ExtDataFig.4F-H). Ectopic formation of spikelet rows was not observed, but we found multi- floret spikelets as described for *Hvclv1* (Fig.4F,G; ExtDataFig.4I), indicating that HvFCP1 acts with HvCLV1 to regulate SAM, IM, SM and RM determinacy.

We next analysed the roles of *HvCLV1* and *HvFCP1* in regulating meristem growth and determinacy using confocal imaging and cell segmentation followed by computational 3D reconstructions of WT, *Hvclv1-1* and *Hvfcp1-1* IMs from W4.5 to W6.5 (Fig.4H). After W5 and termination of spikelet formation, the sizes of *Hvclv1* and *Hvfcp1* IMs increased more rapidly than WT, due to enhanced cell proliferation (Fig.4I,J; ExtDataFig.5A-D). Sizes of the RM were also increased at all stages in *Hvclv1-1* and *Hvfcp1-1,* compared to WT (Fig.5A,B), while FMs were less affected (Fig.5C). *Hvfcp1* mutant phenotypes were overall milder than those of *Hvclv1*, and loss of *HvFCP1* activity did not enhance the phenotype of *Hvclv1* in the *Hvclv1;Hvfcp1* double mutant (Fig.5D), indicating that other CLE peptides can partially compensate for the loss of HvFCP1 and signal through HvCLV1. To analyse this further, we crossed the *HvCLV1* and *HvFCP1* reporter lines into the *Hvfcp1* and *Hvclv1* mutant backgrounds (Fig.5E-L). Expression of *pHvCLV1:HvCLV1-mVenus* was unaltered in the *Hvfcp1-1* IM (Fig.5E,F), while *HvFCP1* expression was reduced in *Hvclv1-1* (Fig.5I-L). Interestingly, HvCLV1 protein internalization, which indicates receptor turnover upon ligand binding^22^, was still detected in the IM and RM in a *Hvfcp1* background, suggesting that in the absence of HvFCP1 an additional peptide can partially compensate its function (Fig.5G,H).

**Fig. 5:**
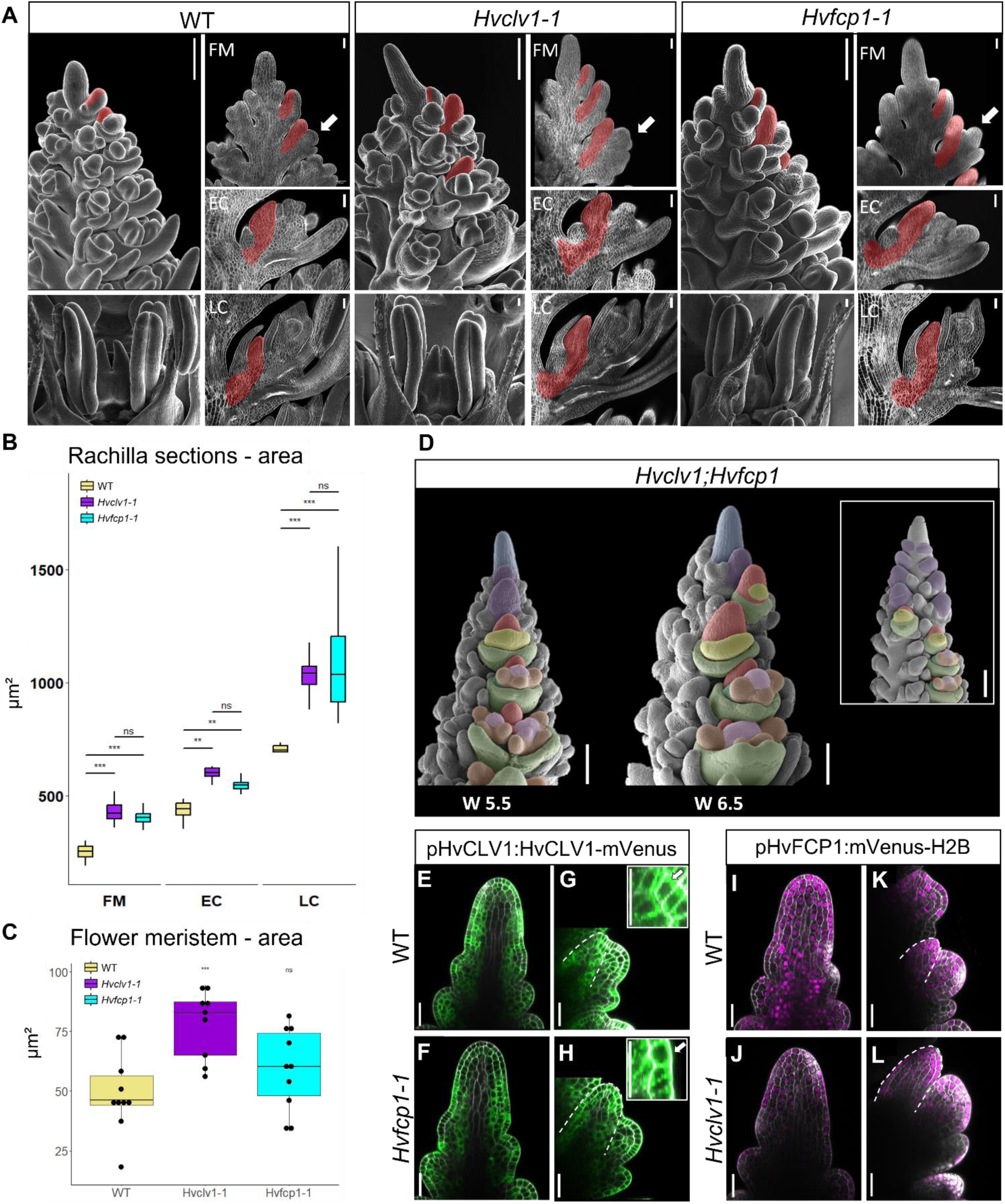
HvCLV1 and HvFCP1 repress RM elongation, *Hvclv1;Hvfcp1* double mutant and reporter lines in the respective mutant backgrounds. **(A)** SEM pictures of WT, *Hvclv1-1* and *Hvfcp1-1* inflorescence meristems at W6.5 used as reference for matching longitudinal sections on their right. The RM in three stages of flower development is highlighted in red. Spikelet with floret meristem (FM), early carpel stage (EC), late carpel stage (LC). White arrows indicate the considered spikelet stage described as FM **(B)** Boxplots displaying rachilla area from central longitudinal sections in WT (yellow), *Hvclv1-1* (purple), *Hvfcp1-1* (cyan). **(C)** Measurements of FM area from SEM frontal pictures of barley inflorescences at W5.5 in WT, *Hvclv1-1* and *Hvfcp1-1*. Dots represents biological replicates and asterixis indicate the significant difference to WT. n=10. **(D)** Inflorescence phenotype of *Hvclv1;Hvfcp1* double mutant at W5.5 and W6.5. On the top right crowned tip phenotype. Color code as described in Fig.1E. **(E-H)** HvCLV1 proteins localization (green) in WT (E,G) and *Hvfcp1-1* (F,H) IM and RM (segmented line) respectively. **(I, L)** *HvFCP1* expression pattern (magenta) respectively in WT and *Hvclv1-1* IM. (R, S) *HvFCP1* expression pattern (magenta) in WT (I,K) and *Hvfcp1-1* (J,L) IM and RM (segmented line) respectively. Scale bars in A: SEM pictures of the inflorescence tip = 200 µm; SEM pictures of flowers and all sections = 50 µm, in D = 200 µm, in E-L = 50 µm.

### HvFCP1 and HvCLV1 control meristematic proliferation through coordination of cell division, auxin signalling and trehalose-6-phosphate

To investigate the common function of HvCLV1 and HvFCP1, we performed RNA-sequencing of WT, *Hvclv1-1* and *Hvfcp1-1* inflorescences at W3.5. A total of 1,208 genes were upregulated and 1,197 downregulated in *Hvclv1* vs WT, while 521 and 258 were upregulated and downregulated in *Hvfcp1* vs WT respectively. Interestingly, 55.2% (288) of the upregulated and 39.9% (103) of the downregulated genes in *Hvfcp1* vs WT were in common with *Hvclv1* vs WT, suggesting a partially shared function of HvFCP1 in the larger gene regulatory network affected by HvCLV1 (ExtDataFig.6A, B).

**Fig. 6:**
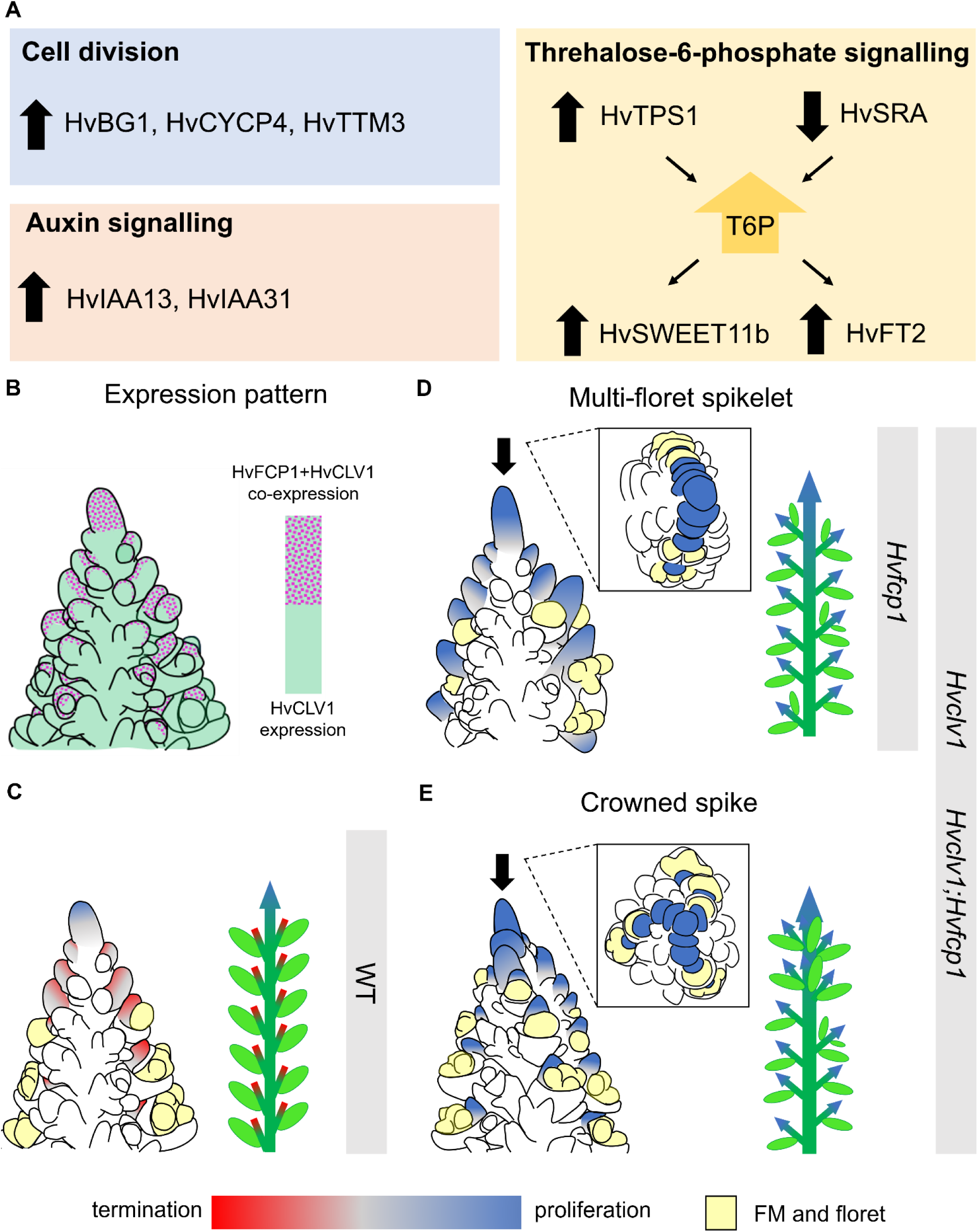
Comparative transcriptome analysis of common gene regulation in *Hvclv1* and *Hvfcp1* vs WT, and schematic summary. **(A)** Commonly regulated genes in *Hvclv1* vs WT and *Hvfcp1* vs WT from RNA-sequencing results. Black arrows pointing upward indicate commonly upregulated genes, while arrows pointing downward indicate commonly downregulated genes. **(B)** Schematic representation of HvCLV1 (green) and *HvFCP1* (magenta dots) expression patterns in barley inflorescence at W5.5. **(C-E)** Schematic representation of barley inflorescences at W5.5 (left) and mature spikes (right) in WT (C), *Hvfcp1*, *Hvclv1* and *Hvclv1;Hvfcp1* (D,E). Grey bars indicate the observed spike phenotypes (multi-floret spikelets and crowned spikes) in the respective genetic backgrounds. Colour code: IM and RMs are marked in red or blue to indicate meristematic proliferation or termination respectively. FM and floral organs are marked in yellow, main rachis and grains in dark and light green.

Mutation of the HvFCP1/HvCLV1 signalling pathway resulted in an enhanced proliferation of the IM and RM in comparison to WT, which ultimately repressed spikelet formation and promoted inflorescence branching. Within the upregulated genes common to both *Hvclv1* vs WT and *Hvfcp1* vs WT, we found *HvBG1*, ortholog of *Rice Big Grain1* (RBG1), which promotes cell division and auxin accumulation in meristematic and proliferating tissues when overexpressed in rice^23^. Furthermore, upregulation of the P-type cyclin *HvCYCP4-1* and the bicistronic transcript encoding Triphosphate Tunnel Metalloenzyme 3 (HvTTM3) and CELL DIVISION CYCLE PROTEIN26 (HvCDC26), together with upregulation of the auxin response genes *HvIAA13* and *HvIAA31,* indicated a general promotion of cell division and alteration of auxin signalling ^24–26^.

Inflorescence branching was previously associated with increased levels of Threhalose-6- Phospate (T6P). Mutation of the maize gene *RAMOSA3* (*RA3*), encoding a Threhalose-6- Phospate Phosphatase, led to indeterminate growth of inflorescence auxiliary meristems, that produced long branches bearing additional FMs^12^. Moreover, studies in Arabidopsis linked increased levels of T6P in axillary meristems with enhanced shoot branching via *FLOWERING LOCUS T* (*FT*) and upregulation the sucrose transporter *Sugars Will Eventually be Exported Transporters11* (*SWEET11*)^27^.

In both *Hvclv1* vs WT and *Hvfcp1* vs WT, *SISTER OF RAMOSA3* (*HvSRA*), paralogue of the maize *RA3,* was downregulated, and *HvTPS1*, the closest ortholog of the Arabidopsis Threhalose-6-Phospate Synthase1 (*TPS1*), was upregulated, suggesting an impaired T6P metabolism. Consistent with findings in Arabidopsis, the sucrose transporter *HvSWEET11b* and *HvFT2*, a barley paralogue of FT, were upregulated in both mutants in comparison to WT, indicating a general reallocation of sucrose and alteration of SM identity (Fig.6A, ExtDataFig.6C, Suppl.Table2)^28,29^. Additionally, the barley gene *COM2* was downregulated in Hv*clv1* vs WT, suggesting HvCLV1 has a role in the upstream regulation of this transcription factor involved in repression of spike branching.

## Discussion

In this study we characterised the function of *CLAVATA* signalling components in coordinating the activity of different meristem types within the barley inflorescence, and showed that *HvCLV1*, together with *HvFCP1*, regulates IM and RM proliferation and determinacy.

The localised expression of *HvFCP1* overlapped with only part of the broader expression of *HvCLV1* (Fig.6B), and while both *Hvclv1* and *Hvfcp1* mutants developed multi-floret spikelets as a consequence of their indeterminate and enlarged rachilla, only *Hvclv1* developed crowned spikes. Additionally, the overall weaker phenotype of *Hvfcp1*, together with observation of HvCLV1 protein internalization in *Hvfcp1* background and the *Hvclv1;Hvfcp1* double mutant phenotype, suggests that additional CLE peptides could interact with HvCLV1, and partially rescue the *Hvfcp1* mutant phenotype. Altogether, our results point toward a more general role of HvCLV1 in mediating the downstream transmission of signals triggered by specifically expressed CLE peptides, thereby regulating the proliferation of different meristems in response to internal or external signals. Transcriptional analyses of *Hvclv1* and *Hvfcp1* highlighted a shared regulatory network between HvCLV1 and HvFCP1, directly or indirectly controlling the expression of genes involved in cell division, auxin signalling and T6P metabolism. Therefore, changes in the proliferation and development of meristems leading to inflorescence growth not only affected inflorescence architecture, but also the overall plant architecture. An enhanced activity of the RM came together with upregulation of *HvTPS1* and downregulation of *HvSRA*, which likely results in accumulation of T6P, previously linked with enhanced branching in Arabidopsis, pea, barley and maize^12,27,30,31^. Increased T6P levels were shown to lead to reorganization of sugar transport by transcriptional regulation of *SWEET* genes and upregulation of *FT*-related genes, which are directly involved in spikelet identity and flowering time^27^. *HvFT2* overexpression in barley consistently resulted in early flowering plants with reduced formation of spikelet primordia, similar to the phenotype observed in *Hvclv1* and *Hvfcp1*^32^.

Meristem homeostasis is controlled redundantly by different receptors and peptides. In maize, rice, and Arabidopsis, parallel and antagonistic pathways control IM shape and maintenance^16,33,34^. The relatively mild phenotype of *Hvclv1*, in comparison to the phenotypes described for *td1* in maize or *clv1* in Arabidopsis, is probably the result of partial compensation by additional CLV-related receptors acting in parallel^19,35^. Combining *Hvclv1* with mutations in other *HvBAM* genes from the same clade might further enhance the *Hvclv1* phenotype.

Our study shows how the *CLV* signalling pathway coordinates the determinacy and growth of diverse meristems of barley spikes, and that regionally expressed CLE peptides differentially regulate the proliferation of specific meristems. Here we note an underexplored opportunity to redesign and optimize barley inflorescence architecture by manipulating the regulation of distinct meristem activities. The large diversity of inflorescence architectures that already evolved in grasses indicates that the underlying genetic networks offer a vast, yet hidden potential to encode a much wider morpho-space than what is realised in our current cereal varieties.

## Material and methods

### Phylogenetic analysis

The CLV1 clade was identified by phylogenetic reconstruction of protein kinase superfamily of four monocots (*Oryza sativa*, *Triticum turgidum*, *Zea mays*, and *Hordeum vulgare*), and two dicot species (*Arabidopsis thaliana*, *Solanum lycopersicum*). The proteomes of these species were downloaded from EnsemblePlants (https://plants.ensembl.org/index.html). To identify all the protein kinase domain containing proteins in the selected species’ proteomes, we conducted HMMscan using HMMER V 3.2 (http://hmmer.org)^36^. HMM profiles of the protein protein kinase domain (Pfam 10.0) were downloaded from InterPro^37^ to carry out the HMM matching. Based on the HMMscan result an E-value threshold of < 1e -10 was imposed to identify the protein kinase domains (PF00069) in the given protein sequences. Protein kinase domains were extracted from all the protein sequences using custom Python scripts and were subjected to HMMalign for protein kinase domain alignment^36^. The multiple sequence alignment was used to construct a Neighbor-joining tree (using JTT+CAT model and default parameters) using the FastTree package^38^. Based on the constructed protein kinase domain family tree we identified the CLV1 clade. Next, we extracted the protein kinase domain sequences from the operational taxonomic units of the selected CLV1 clade to further examine the phylogenetic relationships of kinase domains of CLV1 clade. We constructed the receptor-like kinase phylogeny using RAxML (using random seed for tree initiation and non-parametric bootstrapping) for 1000 bootstrap replicates^39^. This was completed using automated model selection criteria to select the best evolutionary model that fits the dataset.

### Plant material and growth conditions

All barley plants used in this study were *cv.* Golden Promise Fast^40^ and were grown in soil (Einheitserde ED73, Einheitserde Werkverband e.V., with 7% sand and 4 g/L Osmocote Exact Hi.End 3-4M, 4th generation, ICL Group Ltd.) under long day (LD) conditions with 16 hours light at 20°C and 8 hours dark at 16 °C. Plants used for microscopy were grown in QuickPot 96T trays (HerkuPlast Kubern GmbH) in a climate chamber, while the plant phenotype was described in plants growing in larger pots (diameter 16.5 cm, height 13 cm) in a greenhouse under the same growing conditions but with temperatures that slightly varied between seasons. Grains where always pregerminated in Petri dishes with water at 4°C for 3 days before being sowed in soil.

### Plasmids construction and plant transformation

The pHvCLV1:HvCLV1-mVenus plasmid was constructed by PCR amplification of a 2,826 bp fragment upstream of the start codon of HvCLV1 (HORVU.MOREX.r3.7HG0747230) as putative regulatory sequence from Morex genomic DNA (gDNA) and cloned by restriction and ligation via a AscI site into a modified pMDC99^41^. The HvCLV1 coding region without stop codon (3,573 bp) was amplified from Morex gDNA and inserted downstream of the promoter by Gateway cloning (Invitrogen). A C-terminal mVENUS was integrated downstream of the gateway site by restriction and ligation via PacI and SpeI (Suppl.table1). The pHvFCP1:VENUS-H2B construct was cloned by amplifying the regulatory sequence including 2,034 bp upstream of the start codon of HvFCP1 (HORVU.MOREX.r3.2HG0174890) and inserted by Gateway cloning (Invitrogen) into the modified pMDC99^41^. This modified pMDC99 contained the gateway cassette, the coding sequence of VENUS and a T3A terminator, which were inserted by restriction via AscI and SacI from pAB114^42^. Furthermore, it contains the coding sequence of Arabidopsis HISTONE H2B (AT5G22880) at the C terminus of VENUS for nuclear localization, inserted via restriction and ligation at a PacI restriction site (Suppl.table 1). Both pHvCLV1:HvCLV1-mVenus and pHvFCP1:VENUS-H2B constructs were first transformed in the barley cultivar Golden Promise^43^ and then crossed into Golden Promise Fast. *Hvclv1* and *Hvfcp1* mutant alleles were generated by CRISPR-Cas9 genome editing. Plasmids were constructed using the vector system and following the established protocol^44^. The HvCLV1 gene was targeted by a single 20bp sgRNA 53bp after the coding sequence started, while two 20bp sgRNAs were cloned to target HvFCP1 298 and 542 bp after the start codon. All the sgRNAs were designed using E-CRISP software^45^ and single sgRNA strands were hybridized and cloned into the shuttle vectors pMGE625 or pMGE627 by a BpiI cut/ligation reaction. A second cut/ligation reaction (BsaI) was used to transfer the gRNA transformation units (TUs) to the recipient vector pMGE599^44^. The final vector targeting HvCLV1 was transformed in Golden promise Fast via embryo transformation^46^, while the vector targeting HvFCP1 was transformed in Golden Promise Fast via embryo transformation, but using the transformation protocol by Hensel et al. (2009)^47^. Successful insertion of the transformation vector into the genome was tested by PCR (Suppl.table1) on M0 plants. The Cas9 protein was removed by segregation in M1 plants, and homozygous mutations of HvCLV1 and HvFCP1 were identified in M2 plants by amplification of genomic sequences targeted by the sgRNAs and subsequent Sanger sequencing. (Suppl.table1).

### smRNAfish

Barley inflorescences at W3.5 fixed in 4% PFA were embedded in paraplast (Leica Paraplast X-tra) and tissue sections (10 µm) were placed within the capture areas on Resolve Bioscience slides and incubated on a hot plate for at least 20min at 60°C to attach the samples to the slides. Slides were deparaffinized, permeabilized, acetylated and re-fixed. Sections were mounted with a few drops of SlowFade-Gold Antifade reagent (Invitrogen) and covered with a coverslip to prevent damage during shipment to Resolve BioSciences (Germany).

### Plant phenotyping

WT, *Hvclv1* and *Hvfcp1* plants were phenotyped at the end of their life cycle, when completely dried. Plant measurements and percentage of crowned spikes and multi-grains were performed in all the tillers with no distinction between main stem and lateral branches, since the spike phenotypes raised with the same probability in both main and lateral tillers. Three replicates were performed and three plants per replicate were phenotyped.

### Sample preparation, microscopy and image processing

Barley SAMs and inflorescences were collected by manual removal of all the surrounding leaves. Smaller leaves were dissected under a stereo microscope using a 1.5 mm blade scalpel. Fresh inflorescences were directly imaged for stereo microscope pictures using a Nikon SMZ25 stereo microscope with Nikon DS-Fi2 camera. For confocal imaging, fresh barley inflorescences were stuck on their side on a double-sided adhesive tape on an objective slide, stained with 4′,6-diamidin-2-phenylindol (DAPI 1 µg/mL) for 3 minutes, washed three times with water and subsequently covered by with a cover slide before being placed under the microscope. Confocal imaging was performed using Zeiss LSM780 and Zeiss LSM880 with a EC PlnN 10x/0.3, Plan-Apochromat 20x/0.8 or Plan-Apochromat 40x/1 objectives. SEM pictures were obtained by direct imaging of fresh inflorescences or by imaging epoxy replicates of barley inflorescences. At first a negative imprint of the inflorescence was created by mixing the two-component vinyl polysiloxane impression material (Express™ 2Ultra Light Body Quick, 3M ESPE) and pushing the dissected inflorescence into the impression material, which polymerizes a few minutes after having been mixed. After complete polymerization of the negative print, the plant material was removed, and the negative print was filled with epoxy resin. After over-night polymerization, inflorescence replicates were coated with gold using an Agar Sputter Coater and imaged with a Zeiss SUPRA 55VP SEM.

### Peptide treatment

Barley WT and *Hvclv1-1* embryos were dissected at 10 days after pollination, when the SAM was exposed, and cultured on gel media. The medium was prepared by mixing 4.4 g/L of MS medium, 2% sucrose, and 500 ul/L of iron chelate. The pH was adjusted to 6.0 before the addition of 1.5 g/L Gelrite. The medium was then autoclaved. Before being poured in cm square plates, after the medium cooled down, 1/1000 v/v ratio of vitamin mix was added. Treated embryos were then grown in medium with HvFCP1 synthetic peptide (REVPTGPDPIHH, by peptides&elephants GmbH) dissolved in 50ul DMSO reaching a final peptide concentration of 30 µM, while control embryos were grown in medium with 50 µl DMSO. Plates with embryos were grown in a Phyto-cabinet for 30 days at 24°C under long day conditions. After 30 days, when young seedlings developed, vegetative meristems were dissected, fixed in 4% PFA overnight, washed three times with water and incubated for one week in ClearSee solution^48^. Pictures of the cleared meristems were taken under the Zeiss Axioskop 2 light microscope with AxioCam HRc camera. Images were then analysed in Fiji. SAM width was measured by the length of a horizontal line drawn across the SAM base, just on top of the last visible spikelet primordium. Meristem height was calculated as the distance between the SAM tip and the centre of the horizontal line defining the base of the SAM. Three replicates of this experiment were performed and forty embryos were plated for each replicate, even though not all the embryos germinated.

### IM 3D reconstruction and Rachilla vibratome sections

IMs at different W stages were fixed in 4% PFA overnight, then washed three times with water and cleared in ClearSee solution for at least two weeks ^48^. One day before imaging, 1/1000 v/v of SR 2200 stain by Renaissance Chemicals was added to the ClearSee solution and the cell wall was stained. After three washing steps in 1X PBS, barley inflorescences at different stages were glued on the bottom of a small petri dish with a drop of super glue and covered in 1X PBS. The petri dish was placed under Zeiss LSM900 confocal microscope and a z-stack of the submerged IM was imaged from the top with a 20x/0.5 water dipping objective. The 3D reconstruction was performed by loading the IM z-stacks in MorphoGraphX 2.0 and analysed accordingly to the protocol ^49^. Rachilla central longitudinal sections were obtained by following the same procedure described above for the 3D reconstruction. The fixed, cleared inflorescences at W6.5 were embedded in 6% agarose in Disposable Base Molds (epredia) and 50 µm sections were obtained using a Leica VT1000S vibratome. The sections were stained in a Petri dish in 1X PBS with 1/100 v/v concentration of SR 2200 stain for a few minutes, placed on an objective slide with 1X PBS, and covered with a cover slide. Sections of 10 inflorescences for each genotype were then imaged under an LSM880 confocal microscope.

### RNA-sequencing

To detect gene expression changes in *Hvclv1* and *Hvfcp1* inflorescence in comparison to WT, we collected inflorescences of WT, *Hvclv1-1*, and *Hvfcp1-1* at W3.5 for RNA-sequencing. Each replicate contained 40 pooled inflorescences from the main shoot of individual plants. All samples were collected manually under a stereo microscope without surrounding leaves. A total of three biological replicates of each genotype were used for RNA-sequencing. Total RNA was extracted from inflorescences using the Direct-zol^TM^ RNA, Miniprep Plus following the manufacturer’s instructions and digested with DNase I (ZYMO RESEARCH). RNA samples passing a cutoff of RNA Integrity Number (RIN) ≥ 8 were used for mRNA library preparation using poly-A enrichment method. Sequencing was performed on Illumina Novaseq 6000 sequencing platform (PE150), and at least 6G of clean reads data per sample were generated by Biomarker Technologies (BMK) GmbH. To quantify transcripts, all clean reads were mapped to the Morex reference Version 3^50^ using Salmon (v. 0.14.1)^51^. We kept transcripts with a minimum of 1 CPM (counts per million) in at least three samples. Analyses were conducted on 22307 expressed genes. To identify differentially expressed genes (DEGs) within Hvclv1 vs WT and Hvfcp1 vs WT, a pairwise comparisons was conducted using the count-based Fisher’s Exact Test in R package ‘EdgeR’ (v3.32.1)^52^. The FDR of each gene was adjusted by the Benjamini-Hochberg (BH) procedure, thus the gene with BH.FDR<0.05 and log_2_FC ≤ -0.5 or log_2_FC ≥ 0.5 was referred to as downregulated or upregulated gene. The heatmap of gene expression (ExtDataFig.6B) was generated on all the differently expressed genes in *Hvclv1* vs WT and *Hvfcp1* vs WT with -log_10_(TPM + 1) values using ‘ThreeDRNAseq’ R package^53^.

### Quantification and statistical analysis

All the statistical tests were performed using R Studio (RStudio Team 2022). A 2-tailed, unpaired Student’s t-test (function t_test from the package rstatix, v0.7.2) was used to determine the significance between two group means, with a P-value cutoff at ≤ 0.05. Significant difference between more than two groups was determined using a one-way ANOVA (function aov from package stats, v3.6.2) and a subsequent Pairwise t-test (function pairwise.t.test from package stats, v3.6.2), P-value cutoff at ≤ 0.05. Symbols: ns= p-value > 0.05, * = p-value <0.05, ** = p-value <0.01, *** = p-value < 0.001.

## Supporting information

Supplementary informations

## Acknowledgements

We thank Edelgard Wendeler for technical support with barley transformation (MPIPZ), Meik Thiele for providing help with the statistical analysis, Karine Gustavo Pinto for help with plant phenotyping, and Vicky Howe for proofreading. Work in R.S. and M.v.K.S. labs was supported by the DFG through CEPLAS (EXC2048), CSCS (FOR5235), and NEXT-PLANT (IRTG2466).

