## Extended Data Figures for "CLAVATA signalling shapes barley inflorescence architecture by controlling activity and determinacy of shoot apical and rachilla meristems"

### Extended Data Fig.1

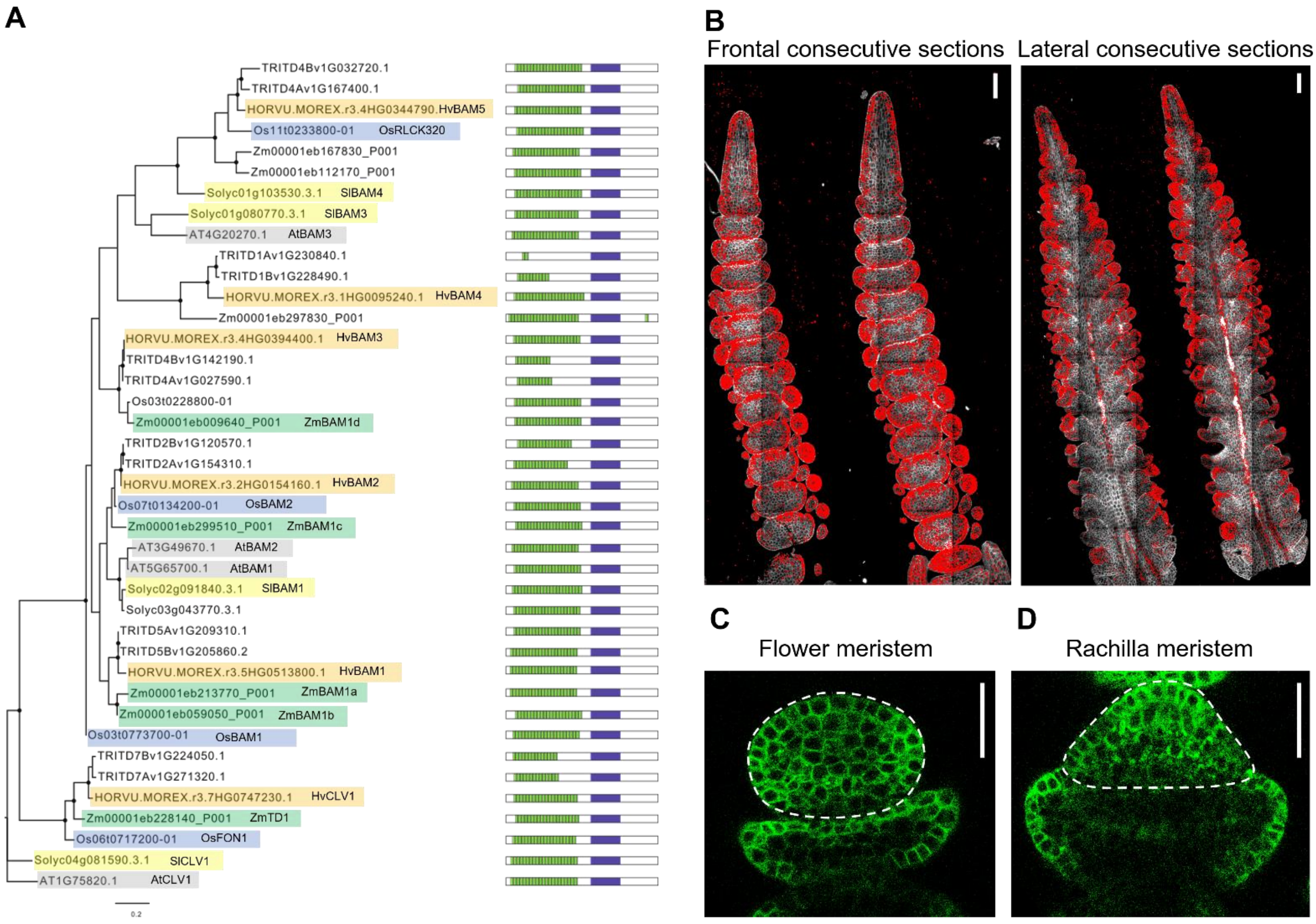


**ExtDataFig1: HvCLV1 clade, expression pattern and protein localization**

**(A)** Maximum likelihood tree of the CLV1 clade based on protein kinase domains. Black dots indicate nodes with bootstrap value higher than 80. Genes identifiers and names of already described genes are highlighted in different colours based on the species [*Arabidopsis thaliana* (grey), *Solanum lycopersicum* (yellow), *Zea mays* (green), *Oryza sativa japonica* (blue) and *Hordeum vulgare* (orange)] are coupled with a schematic representation of the protein structure. Kinase domain as purple rectangle and LRRs as green rectangles. **(B)** Results from smRNAfish (Molecular CartographyTM, Resolve Biosciences) experiment. The fluorescent signal in red shows the localization of HvCLV1 transcripts in dorsal and lateral sections of the barley inflorescence at W3.5. Scalebar: 100 µm **(C,D)** HvCLV1 protein localization (green) in dorsal sections of the floret meristem and rachilla meristem respectively. Scalebar: 50 µm.

### Extended Data Fig.2

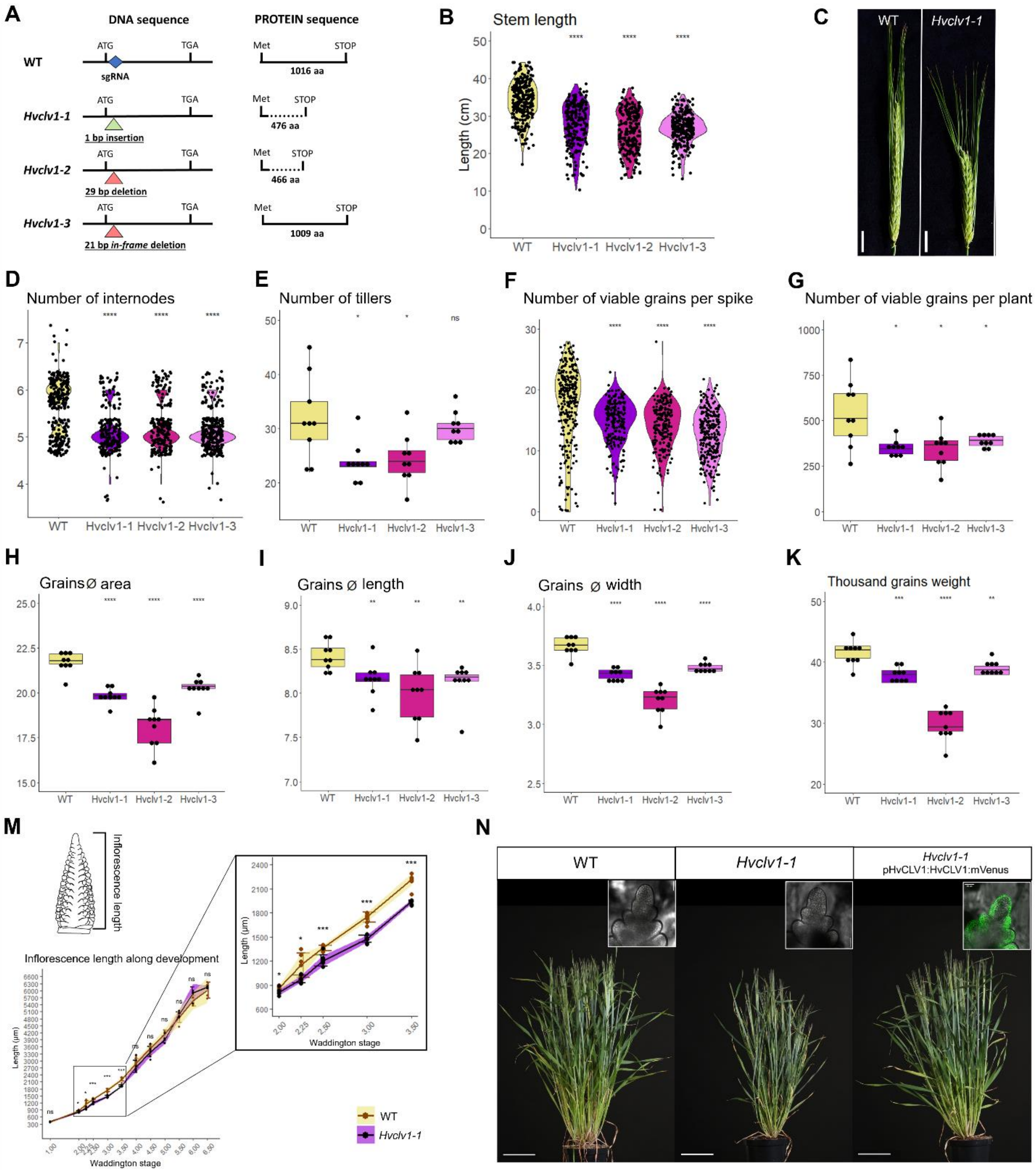


**ExtDataFig2: *Hvclv1* mutant alleles phenotype and complementation with HvCLV1 reporter line**

**(A)** Schematic representation of the HvCLV1 DNA and protein sequence in WT and *Hvclv1-1, Hvclv1-2, Hvclv1-3* mutant alleles. The blue square indicates the region targeted by the sgRNA, the triangles indicate the position of the selected mutation (insertions in green, deletions in red). The dotted lines indicate a different protein sequence to WT. **(B-G)** Plant and spike phenotype. Measurements of stem length (B), spike phenotype of WT and *Hvclv1-1* 60 DAS. Scale bar: 1.5 cm (C), number of internodes (D), number of tillers (E), number of viable grains per spike (F) and per plant (G). Dots indicate single measurement performed in nine mature plants, and asterisks indicate the significant difference in comparison to WT. **(H-K)** Grain phenotype and weight: measurements of grain area (H), grain length (I), grain width (J) and Thousand grains weight (TGW) (K). Dots indicate the average of value of measurements taken in 150 mature grains from nine different plants, asterisks indicate the significant difference in comparison to WT. **(M)** Inflorescence length of WT (yellow) and *Hvclv1-1* (purple). Measurements were taken from W1 to W6.5. Zoom-in plot between W2 and W3.5 on the right. Dots represent single measurements; error bars represent standard deviation and the colored ribbons the interval of confidence. Asterisks indicate the significant difference to WT for each W. **(N)** Complementation of *Hvclv1-1* plant phenotype with HvCLV1 translational reporter line. From left to right WT plant, *Hvclv1-1* and *Hvclv1-1* carrying pHvCLV1:HvCLV1:mVenus plasmid (60 DAS). Pictures on the top right shows HvCLV1:mVenus expression (green) in the IM in the relative genetic background. Scale bar: 10 cm (plants), 50 µm (IMs).

### Extended Data Fig.3

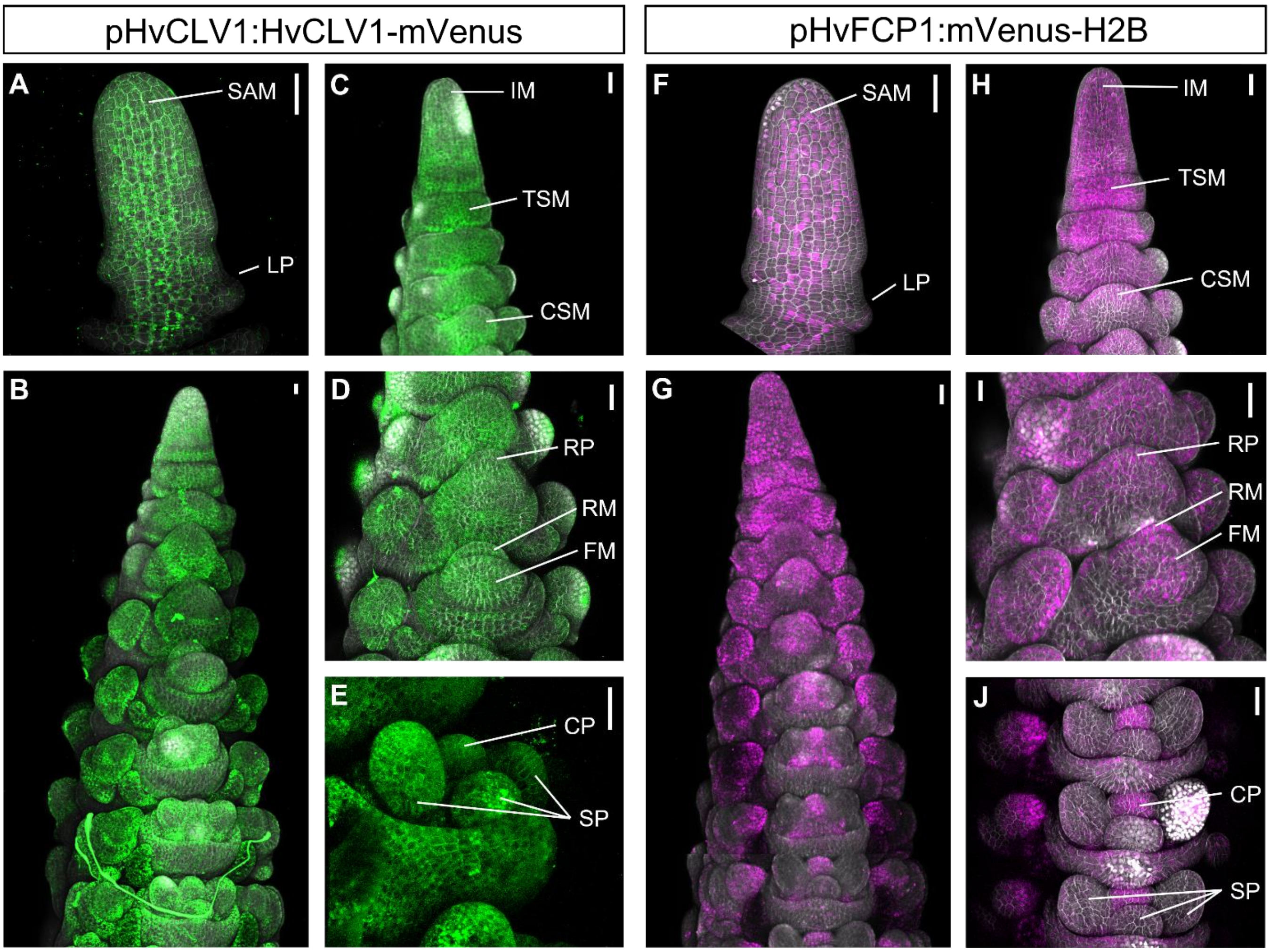


**ExtDataFig3: Comparison between HvCLV1 protein localization and HvFCP1 promoter activity in barley inflorescence**

**(A-E)** HvCLV1 translational reporter line in barely inflorescence at W1 (A) and at W3.5 (B). Close-ups on HvCLV1 protein localization in IM, TSM and CSM (C), RP, RM and FM (D) and CP, SP (E). HvCLV1 proteins in green, DAPI-stained cell wall in grey. **(F-J)** HvFCP1 transcriptional reporter line in barely inflorescence at W1 (F) and at W3.5 (G). Close-ups on HvFCP1 expression in IM, TSM and CSM (H), RP, RM and FM (I) and CP, SP (J). Nuclear localized HvFCP1 expression in magenta, DAPI-stained cell wall in grey. Scalebar: 50 µm.

### Extended Data Fig.4

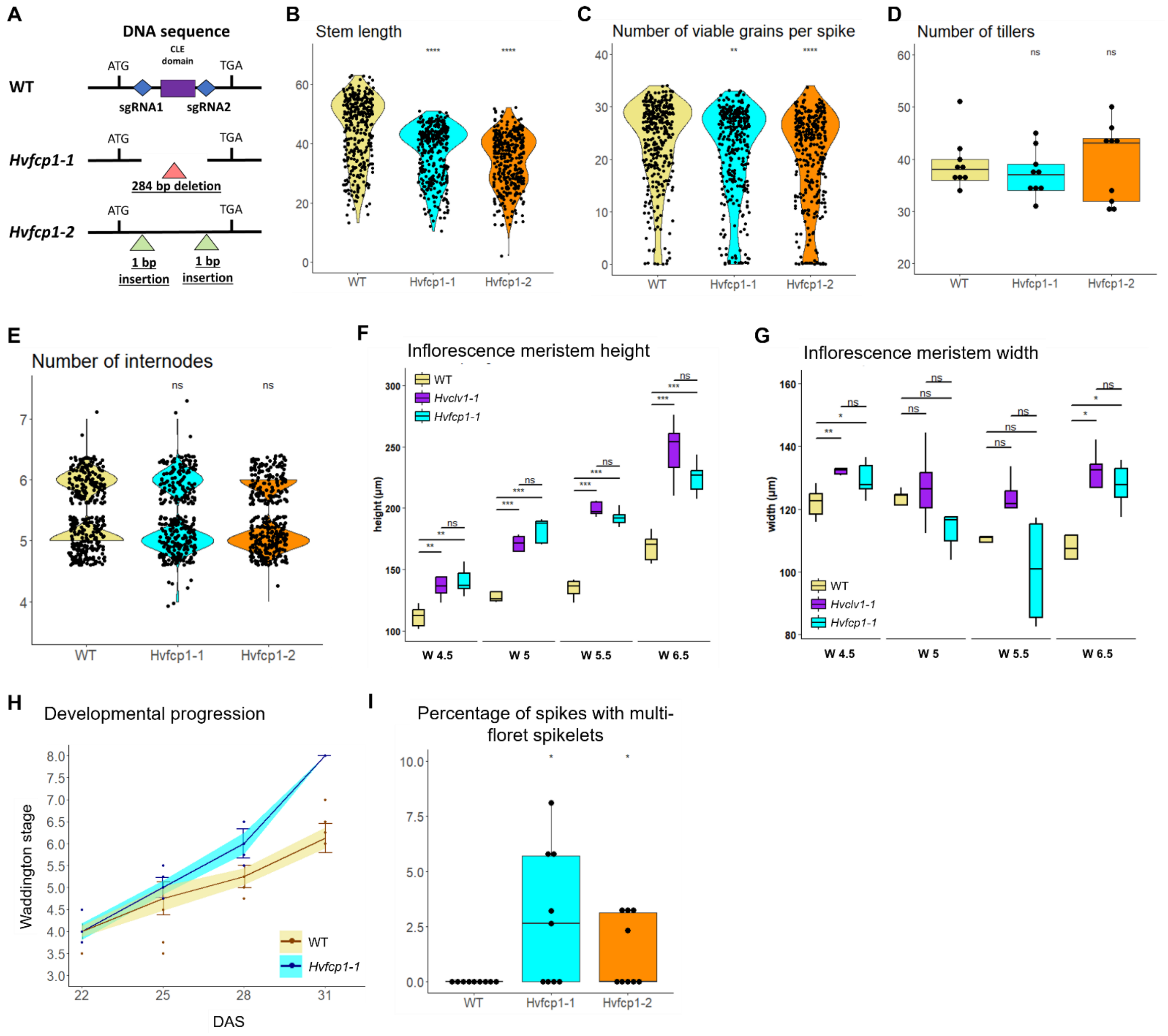


**ExtDataFig4: *Hvfcp1* mutant alleles phenotype and IM shape**

**(A)** Schematic representation of the HvFCP1 DNA sequence in WT and *Hvfcp1-1* and *Hvfcp1-2* mutant alleles. The blue squares indicate the regions targeted by the sgRNAs, the purple rectangle indicates the position of the HvFCP1 CLE domain, and triangles indicate the position of the selected mutation (insertions in green, deletions in red). **(B-E)** Plant phenotype: measurements of stem length (B), number of viable grains per spike (C), number of tillers (D), number of internodes (E). Dots indicate single measurement performed in nine mature plants and asterisks indicate the significant difference in comparison to WT. **(F, G)** Developmental progression and number of spikelet primordia respectively. WT in yellow, *Hvclv1-1* in purple and *Hvfcp1-1* in cyan. Dots represent single measurements; error bars represent standard deviation and the colored ribbons the interval of confidence. **(H)** Developmental progression in WT (yellow) and *Hvfcp1-1* (cyan). Dots represents single measurements; error bars represent standard deviation and the coloured ribbon the interval of confidence. n=10. **(I)** Percentage of spikes with multi-floret spikelets in WT plants and *Hvfcp1* mutant alleles. Dots represent the percentage per plant and asterixis indicate the significant difference to WT. n=9.

### Extended Data Fig.5

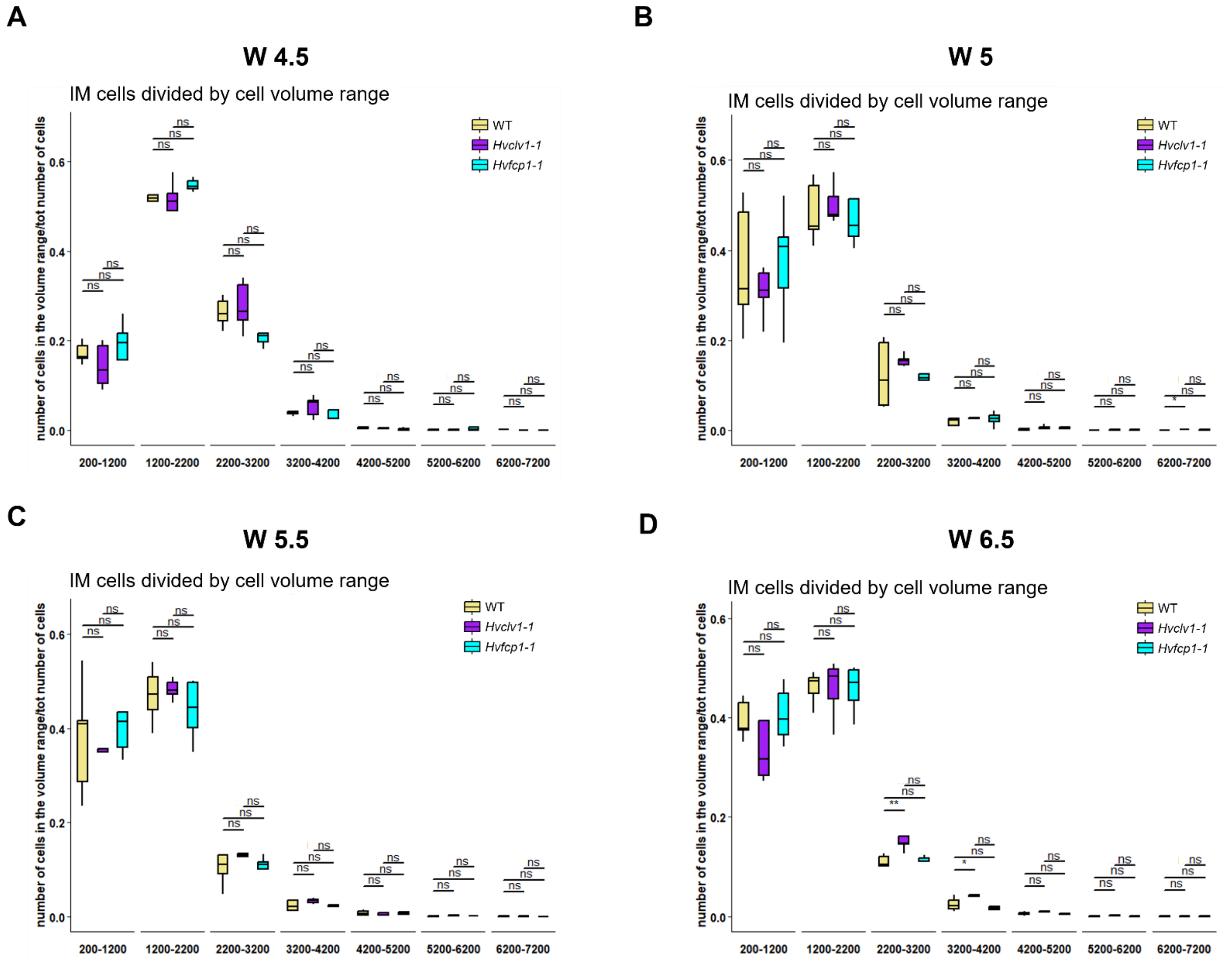


**ExtDataFig5: cell volume is unaffected in *Hvclv1* and *Hvfcp1* IM**

**(A, B, C, D)** Quantification of the Cell Volume Ratio (CVR) in WT (yellow), *Hvclv1-1* (purple) and *Hvfcp1-1* (cyan) in barley inflorescences at W4.5, W5, W6 and W6.5. In the y-axis CVR (ratio between number of IM cells in a specific volume range and total number of cells of the IM), in the x-axis cell volume ranges in cubic micrometers. Boxplots shows the CVR of 5 IMs per Waddington stage.

### Extended Data Fig.6

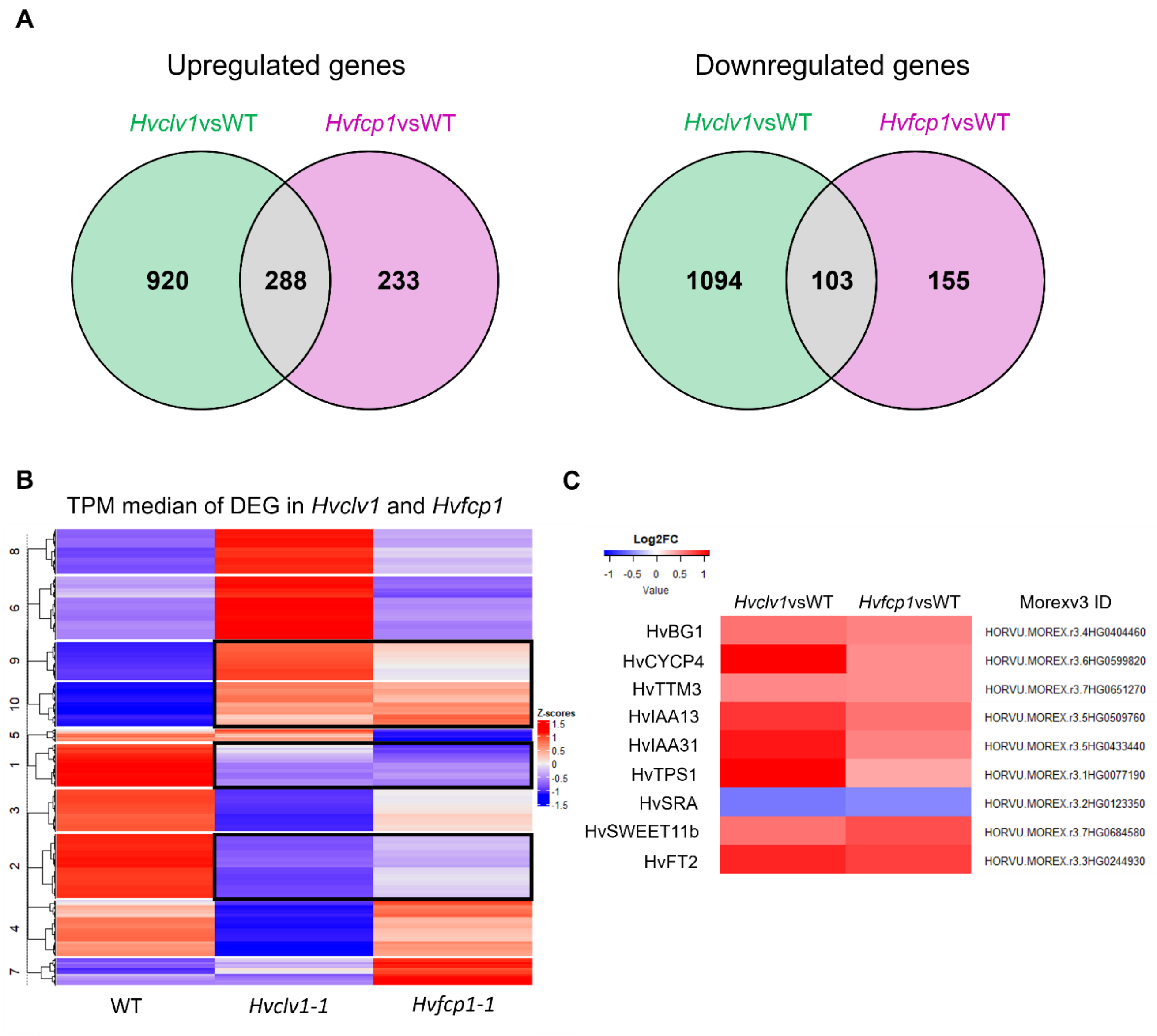


**ExtDataFig6: RNA-seq in *Hvclv1*vsWT and *Hvfcp1*vsWT revealed a common gene regulatory network**

**(A)** Venn diagrams illustrating the number of upregulated (Log2FC > 0.5) and downregulated (Log2FC < -0.5) genes in *Hvclv1*vsWT (green) and *Hvfcp1*vsWT (magenta). Commonly regulated genes in grey. **(B)** Heatmap illustrating the z-score of median transcripts per million (TPM) values for each of the differently expressed genes (DEG) in *Hvclv1* vs WT and *Hvfcp1* vs WT. Clusters on the y-axis group genes with a similar expression trend between genotypes. Black rectangles highlight genes that are similarly regulated in *Hvclv1-1* and *Hvfcp1-1*. (C) Heatmap displaying Log2FC values of genes mentioned in Fig.6A in *Hvclv1*vsWT and *Hvfcp1*vsWT.
