## Supplementary informations for "CLAVATA signalling shapes barley inflorescence architecture by controlling activity and determinacy of shoot apical and rachilla meristems"

### SupplInfo1: *Hvclv1* mutant alleles

#### HvCLV1 (HORVU.MOREX.r3.7HG0747230) Location: Chr7H 618,787,699 - 618,791,569

*Hvclv1-1* carries a 1bp insertion after 70bp that caused a shift in the reading frame and an early stop codon after 476 amino acids (aa), generating a misfolded protein which only shares the first 23aa with the WT sequence (1016aa). *Hvclv1-2* carries a 29bp deletion 41bp after the coding start that caused a shift in the reading frame and an early stop codon after 466 aa. *Hvclv1-3* carries an in-frame deletion of 21 bp after 63bp from the coding start, that allowed the formation of a protein almost identical to the WT apart from 7 missing amino acids (SGSPDRD) in position 22 to 28 in the N-terminal region that disrupt the signal peptide sequence.

<u>Alignment: sgRNA target sequence (blue), insertion (green), deletion (red)</u>

```
HvCLV1          ATGCCGCCACCTCACCTGCTCACCATCCTCCTACCTCTCCTCCTCCTCCTCCCGGCCCCT
Hvclv1-1        ATGCCGCCACCTCACCTGCTCACCATCCTCCTACCTCTCCTCCTCCTCCTCCCGGCCCCT
Hvclv1-2        ATGCCGCCACCTCACCTGCTCACCATCCTCCTACCTCTCCTCC-----------------
Hvclv1-3        ATGCCGCCACCTCACCTGCTCACCATCCTCCTACCTCTCCTCCTCCTCC-----------

                *******************************************

HvCLV1          TCCTCCGGCT-CCCCGGACCGCGACATCTACGCGCTCGCCAAGATCAAGGCCGCCCT…
Hvclv1-1        TCCTCCGGCTTCCCCGGACCGCGACATCTACGCGCTCGCCAAGATCAAGGCCGCCCT…
Hvclv1-2        -------------CCGGACCGCGACATCTACGCGCTCGCCAAGATCAAGGCCGCCCT…
Hvclv1-3        ----------TCCCCGGACCGCGACATCTACGCGCTCGCCAAGATCAAGGCCGCCCT…
                             ********************************************
```

<u>Signal peptide and cleavage position prediction (PredSi)</u>

| | |
|---|---|
| **Matrix:** | Eukarya |
| **Truncation:** | 70 residues |
| **Cleavage position:** | 23 |
| **Score:** | 0.9205 |
| **Secreted protein:** | predicted for secretion |

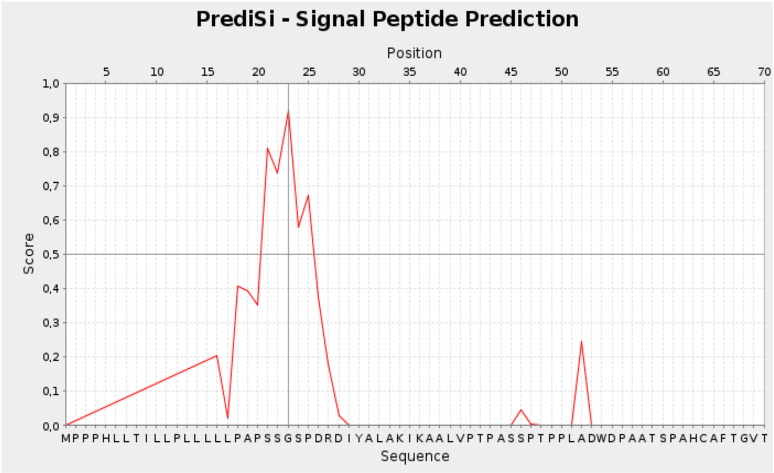


**HvCLV1:**

```
MPPPHLLTILLPLLLLLPAPSSGSPDRDIYALAKIKAALVPTPASSPTPPLADWDPAATSPAHCAFTGVT
|   |    |    |    |    |    |    |    |    |    |    |    |    |    |
1        10        20        30        40        50        60        70
```

HvCLV1 WT protein sequence

MPPPHLLTILLPLLLLLPAPSSGSPDRDIYALAKIKAALVPTPASSPTPPLADWDPAATSPAHCAFTGVTCDA
ATSRVVAINLTALPLHAGTLPPELALLDSLTNLTIAACSLPGRVPAGLPSLPSLRHLNLSNNNLSGPFPAGDG
QTTLYFPSIEVLDCYNNNLSGPLPPFGAAHKAALRYLHLGGNYFSGPIPVAYGDVASLEYLGLNGNALSGRIP
PDLARLGRLRSLYVGYFNQYDGGVPPEFGGLRSLVLLDMSSCNLTGPIPPELGKLKNLDTLFLLWNRLSGEIP
PELGELQSLQLLDLSVNDLAGEIPATLAKLTNLRLLNLFRNHLRGGIPGFVADLPDLEVLQLWENNLTGSLPP
GLGRNGRLRNLDVTTNHLTGTVPPDLCAGGRLEMLVLMDNAFFGPIPESLGACKTLVRVRLSKNFLSGAVPAG
LFDLPQANMLELTDNLLTGGLPDVIGGGKIGMLLLGNNGIGGRIPPAIGNLPALQTLSLESNNFTGELPPEIG
RLRNLSRLNVSGNHLTGAIPEELTRCSSLAAVDVSRNRLTGVIPESITSLKILCTLNVSRNALSGELPTEMSN
MTSLTTLDVSYNALTGDVPMQGQFLVFNESSFVGNPGLCGGPLTGSSNDDACSSSSNHGGGGVLSLRRWDSKK
MLVCLAAVFVSLVAAFLGGRKGCEAWREAARRRSGAWKMTVFQQRPGFSADDVVECLQEDNIIGKGGAGIVYH
GVTRGGGAELAIKRLVGRGVGGDRGFSAEVGTLGRIRHRNIVRLLGFVSNRETNLLLYEYMPNGSLGEMLHGG
KGGHLGWDARARVALEAARGLCYLHHDCAPRIIHRDVKSNNILLDSAFEAHVADFGLAKFLGGAAGASECMSA
IAGSYGYIAPEYAYTLRVDEKSDVYSFGVVLLELITGRRPVGGFGDGVDIVHWVRKATAELPDTAAAVLAVAD
CRLSPEPVPLLVGLYDVAMACVEEASTDRPTMREVVHMLSQPALVAPTAVVDENTARPDDDLILSF*

*Hvclv1* mutant alleles protein sequence

*Hvclv1-1*

MPPPHLLTILLPLLLLLPAPSSGFPGPRHLRARQDQGRPRAHPRILPDAAARRLGPGGDIPSPLRIHRRHMRR
RHLPRRRHQPHRPPAPRRHAAPGARPPRLPNQPHHRRLLPPRPRPRGPPVPAIPPPPQPLQQQPLRPLPRRRR
TDNVVLPVHRGPRLLQQQPLRPAPALRRRAQGRAPLPPPRRELLLRPHPGGLRRRRQPRVPRPQRQRALRQDP
AGPGPAGPAPEPLRRLLQPVRRRRAARVRRAAQPRAARHEQLQPHRPHPARARQAQEPRHALPPLEPIVWRDS
ARAGGAPEPPVAGPVRQRPRRRDTGDPGQAHEPQAAQPVPEPPPRRDTRVRRRPAGPRGAAALGEQPHRQPPA
GTRAQRPAQEPRRHHQPPHRHRAAGPLRGREARDARAHGQRLLRPHPGVAGRVQDAGARPPQQELPQRRRAGR
ALRPAAGQHARAHRQPAHGRPPRRDRRRQDRHAAAGE*

*Hvclv1-2*

MPPPHLLTILLPLLPGPRHLRARQDQGRPRAHPRILPDAAARRLGPGGDIPSPLRIHRRHMRRRHLPRRRHQP
HRPPAPRRHAAPGARPPRLPNQPHHRRLLPPRPRPRGPPVPAIPPPPQPLQQQPLRPLPRRRRTDNVVLPVHR
GPRLLQQQPLRPAPALRRRAQGRAPLPPPRRELLLRPHPGGLRRRRQPRVPRPQRQRALRQDPAGPGPAGPAP
EPLRRLLQPVRRRRAARVRRAAQPRAARHEQLQPHRPHPARARQAQEPRHALPPLEPIVWRDSARAGGAPEPP
VAGPVRQRPRRRDTGDPGQAHEPQAAQPVPEPPPRRDTRVRRRPAGPRGAAALGEQPHRQPPAGTRAQRPAQE
PRRHHQPPHRHRAAGPLRGREARDARAHGQRLLRPHPGVAGRVQDAGARPPQQELPQRRRAGRALRPAAGQHA
RAHRQPAHGRPPRRDRRRQDRHAAAGE*

*Hvclv1-3*

MPPPHLLTILLPLLLLLPAPSIYALAKIKAALVPTPASSPTPPLADWDPAATSPAHCAFTGVTCDAATSRVVA
INLTALPLHAGTLPPELALLDSLTNLTIAACSLPGRVPAGLPSLPSLRHLNLSNNNLSGPFPAGDGQTTLYFP
SIEVLDCYNNNLSGPLPPFGAAHKAALRYLHLGGNYFSGPIPVAYGDVASLEYLGLNGNALSGRIPPDLARLG
RLRSLYVGYFNQYDGGVPPEFGGLRSLVLLDMSSCNLTGPIPPELGKLKNLDTLFLLWNRLSGEIPPELGELQ
SLQLLDLSVNDLAGEIPATLAKLTNLRLLNLFRNHLRGGIPGFVADLPDLEVLQLWENNLTGSLPPGLGRNGR
LRNLDVTTNHLTGTVPPDLCAGGRLEMLVLMDNAFFGPIPESLGACKTLVRVRLSKNFLSGAVPAGLFDLPQA
NMLELTDNLLTGGLPDVIGGGKIGMLLLGNNGIGGRIPPAIGNLPALQTLSLESNNFTGELPPEIGRLRNLSR
LNVSGNHLTGAIPEELTRCSSLAAVDVSRNRLTGVIPESITSLKILCTLNVSRNALSGELPTEMSNMTSLTTL
DVSYNALTGDVPMQGQFLVFNESSFVGNPGLCGGPLTGSSNDDACSSSSNHGGGGVLSLRRWDSKKMLVCLAA
VFVSLVAAFLGGRKGCEAWREAARRRSGAWKMTVFQQRPGFSADDVVECLQEDNIIGKGGAGIVYHGVTRGGG
AELAIKRLVGRGVGGDRGFSAEVGTLGRIRHRNIVRLLGFVSNRETNLLLYEYMPNGSLGEMLHGGKGGHLGW
DARARVALEAARGLCYLHHDCAPRIIHRDVKSNNILLDSAFEAHVADFGLAKFLGGAAGASECMSAIAGSYGY
IAPEYAYTLRVDEKSDVYSFGVVLLELITGRRPVGGFGDGVDIVHWVRKATAELPDTAAAVLAVADCRLSPEP
VPLLVGLYDVAMACVEEASTDRPTMREVVHMLSQPALVAPTAVVDENTARPDDDLILSF*

### SupplInfo2: *Hvclv1-1* growing in semi-field conditions

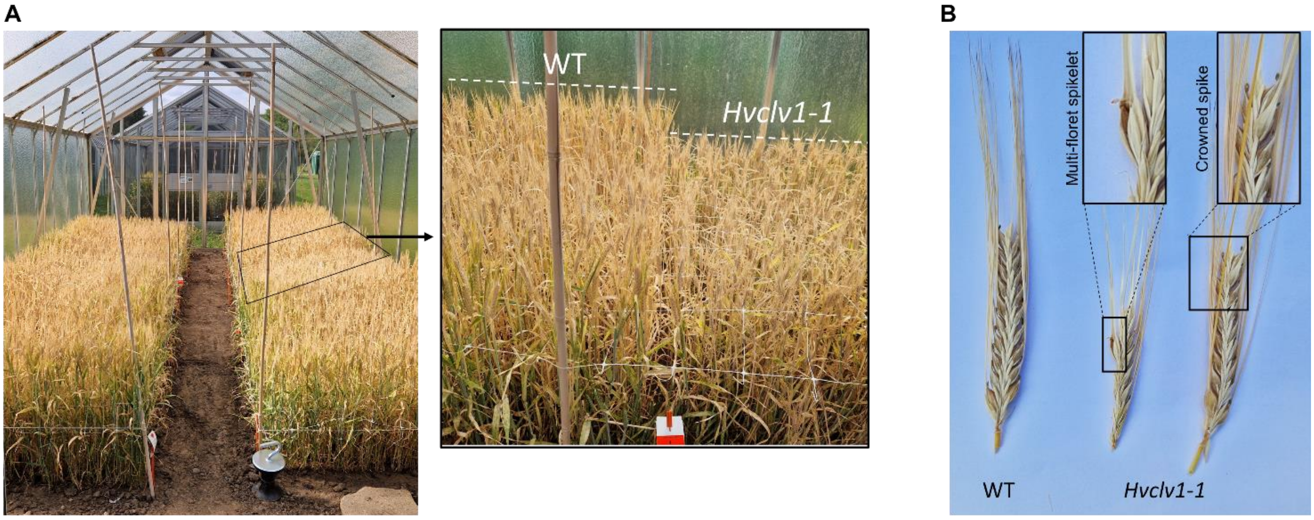


In (A) picture of the semi-field setup and zoom-in in WT and *Hvclv1-1* plants. In every plot twelve rows of 44 grains were sown 11 cm apart (528 grains per 1.3mx1.35m =~300 grains per $m^2$) and grown between March and August 2023. In (B) examples of spikes form WT and *Hvclv1-1* mutant showing multi-floret spikelets and crowned spike. The average plot yield was 851.7 +- 85 grams for WT plants and 466.4 +- 20.4 grams for *Hvclv1-1.* Thousand grain weight (TGW) was 41.9 +- 0.1 for WT plants and 38.0 +- 1.4 for *Hvclv1-1* plants.

### SupplInfo3:

**HvFCP1 (HORVU.MOREX.r3.2HG0174890)** Location: Chr2H 523,068,208 - 523,069,031

We generated two independent knock-out mutant alleles by CRISPR-Cas9, called *Hvfcp1-1* and *Hvfcp1-2*. The 284bp deletion in *Hvfcp1-1* removed part of the first exon and the entire second exon, which normally carries the conserved CLE domain. *Hvfcp1-2*, carries two1bp insertions at the +298bp and +543bp positions. The first insertion caused a shift in the reading frame that altered the entire amino acid sequence of the predicted peptide.

FCP1 CLE domain sequence in barley, maize and rice

```
HORVU.MOREX.r3.2HG0174890(HvFCP1)   REVPTGPDPIHHH
Zm00001d003320(ZmFCP1)              REVPTGPDPIHHH
Os04t0473800(OsFCP1)                REVPTGPDPIHHH
```

sgRNAs target sequences (blue), insertion (green), deletion (red)

```
HvFCP1          ATGGCTCATGCCGCCGACGCGAGGTCGCGCTGCGTCGTCGCGGTGCTCTTCGCCGTAGCC
Hvfcp1-1        ATGGCTCATGCCGCCGACGCGAGGTCGCGCTGCGTCGTCGCGGTGCTCTTCGCCGTAGCC
Hvfcp1-2        ATGGCTCATGCCGCCGACGCGAGGTCGCGCTGCGTCGTCGCGGTGCTCTTCGCCGTAGCC
                ************************************************************

HvFCP1          GTCTTCCTCGCCTGCTTGCCGCCCGCCGCCGCCTCCTCCTCGTCTTCCCGGGCAGGTACG
Hvfcp1-1        GTCTTCCTCGCCTGCTTGCCGCCCGCCGCCGCCTCCTCCTCGTCTTCCCGGGCAGGTACG
Hvfcp1-2        GTCTTCCTCGCCTGCTTGCCGCCCGCCGCCGCCTCCTCCTCGTCTTCCCGGGCAGGTACG
                ************************************************************

HvFCP1          TGCGTCGTCCCGTCCGCCATGCGTTCGTTGCTCTCTACAACCCCCGCCGCAAGGCCACCT
Hvfcp1-1        TGCGTCGTCCCGTCCGCCATGCGTTCGTTGCTCTCTACAACCCCCGCCGCAAGGCCACCT
Hvfcp1-2        TGCGTCGTCCCGTCCGCCATGCGTTCGTTGCTCTCTACAACCCCCGCCGCAAGGCCACCT
```

```
                ************************************************************

HvFCP1          CCCTGGTTCTCGCGCCGCACGGGAATCTCCTGCGCTCTTTGACGCCTTTGTTGGTCATCT
Hvfcp1-1        CCCTGGTTCTCGCGCCGCACGGGAATCTCCTGCGCTCTTTGACGCCTTTGTTGGTCATCT
Hvfcp1-2        CCCTGGTTCTCGCGCCGCACGGGAATCTCCTGCGCTCTTTGACGCCTTTGTTGGTCATCT
                ************************************************************

HvFCP1          CCCTCGCAGCGGCGGCGGCGGCATTGCAACGAGTCGAGATGGCGGCCATGTACACCCCGC
Hvfcp1-1        CCCTCGCAGCGGCGGCGGCGGCATTGCAACGAGTCGAGAT--------------------
Hvfcp1-2        CCCTCGCAGCGGCGGCGGCGGCATTGCAACGAGTCGAGATGGCGGCCATGTACACCCCGC
                ****************************************

HvFCP1          AGGACCTGCAGGAGAAG-CCGGATGTGACCAAGGTACGTACGCGGCCGCCATGTTACGGC
Hvfcp1-1        ------------------------------------------------------------
Hvfcp1-2        AGGACCTGCAGGAGAAAGCCGGATGTGACCAAGGTACGTACGCGGCCGCCATGTTACGGC

HvFCP1          TTCGGGCCGAAGGAAAGGCGGCTCCTTTGGTGGTTTCTTGCTGTCGTGTTTCGAGCTCAT
Hvfcp1-1        ------------------------------------------------------------
Hvfcp1-2        TTCGGGCCGAAGGAAAGGCGGCTCCTTTGGTGGTTTCTTGCTGTCGTGTTTCGAGCTCAT

HvFCP1          GGGGTTTTGATTTTCGATGCGCAGGACGCGGAGGAGGACGTGAGCACGACGGGGTTCGGC
Hvfcp1-1        ------------------------------------------------------------
Hvfcp1-2        GGGGTTTTGATTTTCGATGCGCAGGACGCGGAGGAGGACGTGAGCACGACGGGGTTCGGC

HvFCP1          GCGGAGGAGGAGAGGGAGGTGCCCACCGGGCCGGACCCCATCCACCACCACGGCAGGGGA
Hvfcp1-1        ------------------------------------------------------------
Hvfcp1-2        GCGGAGGAGGAGAGGGAGGTGCCCACCGGGCCGGACCCCATCCACCACCACGGCAGGGGA

HvFCP1          CCCAGG-CGCCGGCAGTCGCCCTGATCGCGCGGCAGGTGGAGGATGCTTCCGTGGGTCG…
Hvfcp1-1        --------------------------CGCGCGGCAGGTGGAGGATGCTTCCGTGGGTCG…
Hvfcp1-2        CCCAGGGCGCCGGCAGTCGCCCTGATCGCGCGGCAGGTGGAGGATGCTTCCGTGGGTCG…
                                          *********************************
```

<u>WT protein sequence</u> (CLE domain highlighted in yellow)

```
MAHAADARSRCVVAVLFAVAVFLACLPPAAASSSSSRAAAAAALQRVEMAAMYTPQDLQE
KPDVTKDAEEDVSTTGFGAEEEREVPTGPDPIHHHGRGPRRRQSP*
```

<u>*Hvfcp1* mutant alleles protein sequence</u>

*Hvfcp1-1*

```
MAHAADARSRCVVAVLFAVAVFLACLPPAAASSSSSRAGTCVVPSAMRSLLSTTPAARPPPWFSRRTGISCAL
*
```

*Hvfcp1-2*

```
MAHAADARSRCVVAVLFAVAVFLACLPPAAASSSSSRAAAAAALQRVEMAAMYTPQDLQEKAGCDQGRGGGRE
HDGVRRGGGEGGAHRAGPHPPPRQGTQGAGSRPDRAAGGGCFRGSVHPA*
```

<u>Alignment</u>

```
HvFCP1          MAHAADARSRCVVAVLFAVAVFLACLPPAAASSSSSRAAAAAALQRVEMAAMYTPQDLQE
Hvfcp1-2        MAHAADARSRCVVAVLFAVAVFLACLPPAAASSSSSRAAAAAALQRVEMAAMYTPQDLQE
Hvfcp1-1        MAHAADARSRCVVAVLFAVAVFLACLPPAAASSSSSRAGTCVVPSAMRSLLSTTPAAR--
                **************************************.:... . :.     **

HvFCP1          KP--DVTKDAEEDVSTTGFGAEEEREVPTGPDPIHHHGRGPRRRQSP-------------
Hvfcp1-2        KAGCDQGRGGGREHDGVRRGGGEGGAHRAGPHPPPRQGTQGAGSRPDRAAGGGCFRGSVH
```

```
Hvfcp1-1        ------------------------------PPPWFSRRTGISCAL---------------
                                              * *   :

HvFCP1          --
Hvfcp1-2        PA
Hvfcp1-1        --
```
